# *Toxoplasma* GRA15 limits parasite growth in IFNγ-activated fibroblasts through TRAF ubiquitin ligases

**DOI:** 10.1101/2020.02.24.963496

**Authors:** Debanjan Mukhopadhyay, Lamba Omar Sangaré, Laurence Braun, Mohamed-Ali Hakimi, Jeroen P.J. Saeij

## Abstract

The protozoan parasite *Toxoplasma gondii* lives inside a vacuole in the host cytoplasm where it is protected from host cytoplasmic innate immune responses. However, IFNγ-dependent cell-autonomous immunity can destroy the vacuole and the parasite inside. *Toxoplasma* strain differences in susceptibility to human IFNγ exist but the *Toxoplasma* effector(s) that determine these differences are unknown. We show that in human primary fibroblasts, the polymorphic *Toxoplasma* secreted effector GRA15 mediates the recruitment of ubiquitin ligases, including TRAF2 and TRAF6, to the vacuole membrane, which enhances recruitment of ubiquitin receptors (p62/NDP52) and ubiquitin-like molecules (LC3B, GABARAP). This ultimately leads to lysosomal degradation of the vacuole. In murine fibroblasts, GRA15-mediated TRAF6 recruitment mediates the recruitment of immunity-related GTPases and destruction of the vacuole. Thus, we have identified how the *Toxoplasma* effector GRA15 affects cell-autonomous immunity in human and murine cells.

## Introduction

*Toxoplasma* is a highly successful obligate intracellular parasite that can establish lifelong chronic infections in a wide range of warm-blooded animals. In humans it causes opportunistic infections in immunosuppressed patients, congenital infections (Hill & Dubey, 2002), and blindness (Pleyer *et al*, 2014). Many different *Toxoplasma* strains exist but in Europe and North-America human infections are dominated by the type I and type II clonal lineages (Howe & Sibley, 1995; Saeij *et al*, 2005). Upon host cell invasion, *Toxoplasma* wraps itself with the host cell plasma membrane, which becomes the nascent parasitophorous vacuole membrane. The vacuole membrane does not fuse with the endo-lysosome system and without immune pressure the vacuole does not acidify thus providing *Toxoplasma* with a niche for replication (Jones *et al*, 1972; Mordue & Sibley, 1997).

Like *Toxoplasma*, many intracellular pathogens reside within a vacuole (pathogen-containing vacuole or PV) in the host cytoplasm (Liehl *et al*, 2015). The PV membrane (PVM) protects these pathogens from detection by host cytosolic pathogen recognition receptor (PRRs). However, the host has developed mechanisms to destroy the PV thereby exposing the pathogen (Liehl *et al*, 2015; Saeij & Frickel, 2017). For example, in mice interferons upregulate the expression of two families of large dynamin-like GTPases: the immunity-related GTPases (IRGs) and the guanylate binding proteins (GBPs) (Howard *et al*, 2011), which mediate the destruction of the PV of *Toxoplasma* and of many gram-negative bacteria (Liehl *et al*, 2015). Once the pathogen is in the cytoplasm, GBPs and IRGs can also mediate the vesiculation of the pathogen itself thereby exposing pathogen associated molecular patterns (PAMPs) to cytosolic PRRs, which can lead to the activation of the inflammasome (Man *et al*, 2017) and the induction of a form of cell death called pyroptosis (Broz & Dixit, 2016). Because host cell death removes the replication niche of intracellular pathogens, this is an efficient way of inhibiting pathogen growth (Krishnamurthy *et al*, 2017).

The mechanism of IRG and GBP recruitment to the *Toxoplasma* PVM and the identity of *Toxoplasma* effectors influencing IRG/GBP recruitment, or their activity, are intense areas of research. These GTPases are normally held inactive on host endomembranes by regulatory ‘GMS motif’ type IRGs (Haldar *et al*, 2013). Initially, ‘pioneer’ effector ‘GKS’ motif type IRGs are recruited to the *Toxoplasma* PVM(Hunn *et al*, 2008), which is largely devoid of regulatory IRGs, where they oligomerize and become activated. What exact signal initiates the recruitment of these pioneer IRGs to the PVM is unclear. In murine cells the initial conjugation of a ubiquitin-like protein (e.g. microtubule-associated protein light chain 3 (LC3) or γ-aminobutyric acid receptor-associated proteins (GABARAPs)) to the PVM was proposed to be the signal that initiates recruitment of ‘pioneer’ IRGs (Choi *et al*, 2014; Sasai *et al*, 2017). PVM recruitment of pioneer IRGs somehow promotes the ubiquitination of the PVM which subsequently leads to the recruitment of the ubiquitin-binding protein p62 (also called sequestosome or SQSTM1) and E3 ubiquitin ligases (e.g. TNF receptor associated factor 6 or TRAF6 and Tripartite motif containing 21 or TRIM21) in a co-dependent manner (Foltz *et al*, 2017; Haldar *et al*, 2015; Lee *et al*, 2015). PVM ubiquitination by these ubiquitin ligases can lead to further recruitment of p62 and TRAF6 thereby creating an amplification loop ensuring the full ubiquitination of the PV. GBPs also get recruited to the PVM and eventually vesiculation of the PVM by GBPs and IRGs expose *Toxoplasma*, which can lead to its destruction by GBPs (Degrandi *et al*, 2013; Kravets *et al*, 2016) and pyro-necrosis of host cells (Zhao *et al*, 2009). To counter the host defense mechanisms, *Toxoplasma* secretes ROP and GRA effector proteins into the host cell from organelles called rhoptries and dense granules, respectively. In both murine and human cells, the type II strain is more susceptible to IFNγ-mediated growth inhibition than the type I strain (Clough *et al*, 2016; Haldar *et al*, 2015; Qin *et al*, 2017; Selleck *et al*, 2015). Resistance of type I strains in murine cells is determined primarily by polymorphic ROP5 and ROP18, which, together with ROP17 and GRA7, cooperatively block IRG and GBP loading on the PVM and subsequent events (Behnke *et al*, 2015; Etheridge *et al*, 2014; Fleckenstein *et al*, 2012; Haldar *et al*, 2015; Niedelman *et al*, 2012; Steinfeldt *et al*, 2010; Khaminets *et al*, 2010). ROP16 and GRA15 also affect GBP loading on the PVM in murine cells through an unknown mechanism (Virreira Winter *et al*, 2011).

IFNγ-stimulated human cells control *Toxoplasma* using diverse mechanisms dependent on the cell type (Krishnamurthy *et al*, 2017). For example, IFNγ-mediated induction of Indoleamine 2,3 dioxygenase (IDO) causes breakdown of L-Tryptophan, for which *Toxoplasma* is auxotrophic, which has been shown to mediate inhibition of parasite growth in HeLa, HAP1, and fibroblast cells (Niedelman *et al*, 2013; Bando *et al*, 2018; Qin *et al*, 2017; Pfefferkorn, 1984; Pfefferkorn *et al*, 1986). In some human cell lines another mechanism for parasite control is ubiquitination of the vacuole, which leads to lysosomal fusion in HUVEC cells, while in HeLa cells an autophagic double membrane forms around the vacuole and parasite growth is stunted without lysosomal fusion (Clough *et al*, 2016; Selleck *et al*, 2015). In certain human cells, GBP1 also seems important for restriction of *Toxoplasma* growth but the mechanism of growth restriction is unclear. In a lung epithelial cell line (A549) GBP1 restricts parasite growth without its recruitment to the PVM (Johnston *et al*, 2016) while in human mesenchymal stem cells (MSC) growth restriction was associated with GBP1 PVM recruitment (Qin *et al*, 2017). Much less is known about what initiates targeting of human immune effectors to the PVM and how this leads to parasite elimination. In contrast to mice, humans lack IFNγ-inducible IRGs, likely explaining why ROP5, ROP18, and ROP17 do not seem to play an important role in conferring protection against IFNγ-mediated growth inhibition in human cells (Clough *et al*, 2016; Niedelman *et al*, 2012; Selleck *et al*, 2015). Currently, no parasite proteins that determine strain differences in susceptibility to IFNγ-mediated cell autonomous immunity in human cells have been identified. A secreted parasite effector, *Toxoplasma* inhibitor of STAT1-dependent transcription (TgIST), which blocks the STAT1 transcriptional response, was recently described but this effector functions upstream of the upregulation of IFNγ-induced toxoplasmacidal mechanisms in both type I and type II strains (Gay *et al*, 2016; Olias *et al*, 2016).

Herein, we report that the PVM-localized *Toxoplasma* GRA15 effector (Rosowski *et al*, 2011) enhances parasite susceptibility to IFNγ in primary human foreskin fibroblast (HFFs) and murine embryonic fibroblasts (MEFs). GRA15 binds several ubiquitin ligases, including TRAF2 and TRAF6, which in HFFs is associated with enhanced recruitment of p62, LC3 and GABARAP to the PVM, enhanced endo-lysosomal fusion, and parasite destruction. Unlike what has been described for murine cells, PVM ubiquitination does not correlate with *Toxoplasma* elimination in IFNγ-stimulated HFFs. The type I RH strain, which does not express a functional GRA15, is much less susceptible to IFNγ-mediated elimination. In MEFs, GRA15 also interacts with TRAF2 and TRAF6, and TRAF6 recruitment leads to enhanced PVM loading with IRGs and GBPs and parasite destruction. Thus, we determined that the *Toxoplasma* effector GRA15 mediates strain differences in susceptibility to cell autonomous immunity in human cells and determined the mechanism by which GRA15 enhances parasite susceptibility to host IFNγ in both human and murine fibroblasts.

## Results

### The polymorphic effector GRA15 enhances *Toxoplasma* susceptibility to IFNγ-mediated growth inhibition in HFFs

To determine if the type I RH and the type II Pru strain differ in susceptibility to IFNγ-mediated growth inhibition in HFFs, we measured the IFNγ-mediated reduction in plaque numbers and plaque area. The relative reduction in the number of plaques formed in IFNγ-stimulated *vs.* unstimulated cells reflects killing of *Toxoplasma,* while the relative reduction in the area of the plaques is a sensitive assay that reflects overall inhibition of parasite growth over multiple lysis cycles (Niedelman *et al*, 2012, 2013). Compared to RH, Pru had a larger IFNγ-induced loss in plaque numbers and plaque area (**Fig. 1a**). We used parasites expressing luciferase to determine IFNγ-mediated growth inhibition 24 h after infection and observed that Pru growth was more inhibited by IFNγ compared to RH growth (**Fig. 1b**). IFNγ stimulation resulted in a decrease in the relative number of vacuoles (**Fig. 1c**) as well as a decrease in the number of parasites per vacuole (**Fig. 1d**), both of which were more pronounced for the Pru strain.

**Figure 1:**
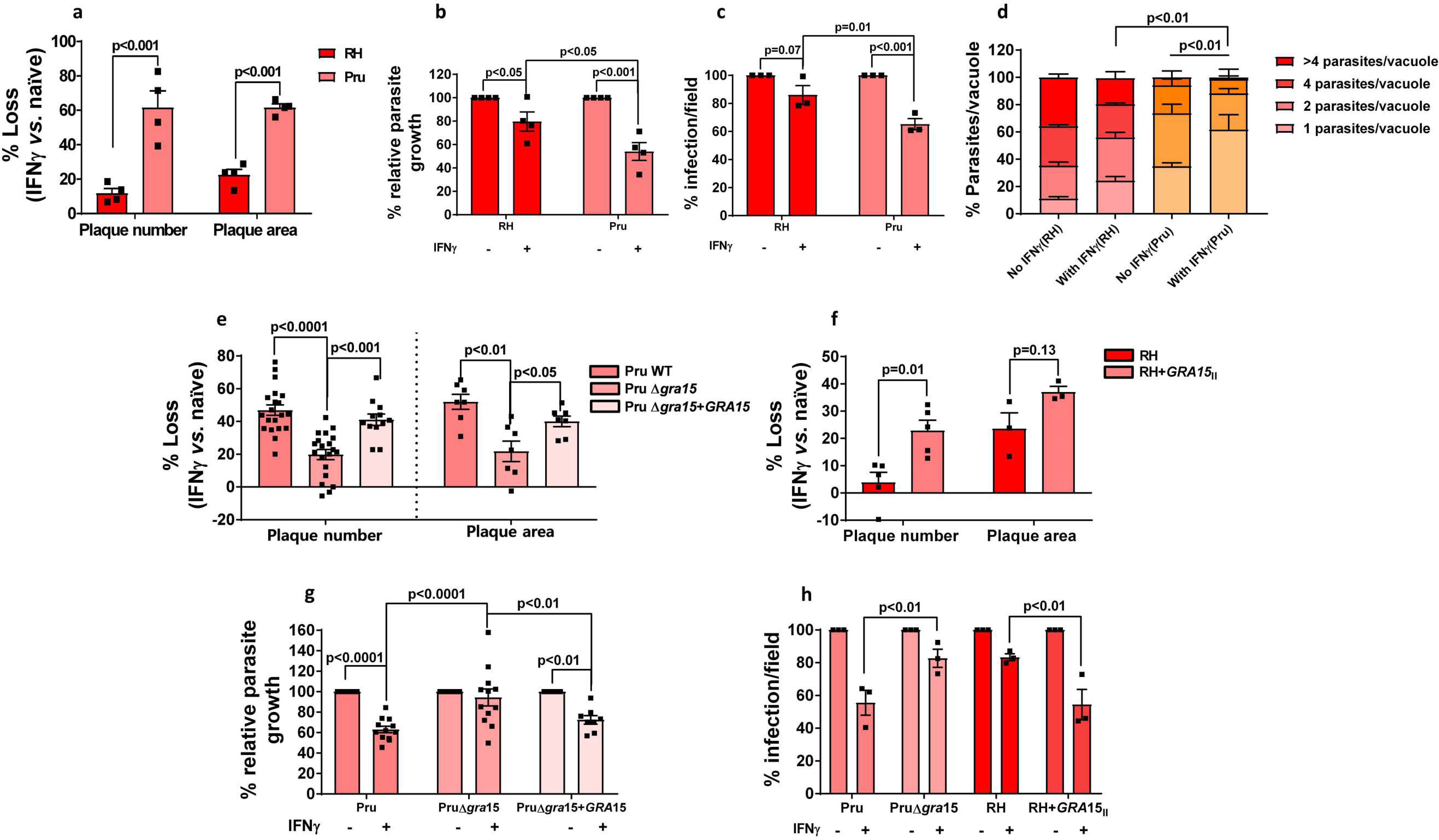
The type II (Pru) *Toxoplasma* strain is more susceptible to IFNγ-mediated growth inhibition in primary human foreskin fibroblasts (HFFs) than the type I RH strain due to presence of the polymorphic effector GRA15. **a)** HFFs were pre-stimulated with IFNγ (10U/mL) for 24 h. Plaque assays were performed for each strain and each condition. Plaque number and area loss were calculated 4 days p.i. for RH and 6 days p.i. for Pru. Assays were performed with RH (*n=4*), Pru (*n=4*). **b)** Relative parasite growth was measured 24 h p.i. in IFNγ-stimulated and unstimulated HFFs by luciferase assay. Growth of each strain in IFNγ-stimulated HFFs is expressed relative to growth in unstimulated HFFs. Experiments were performed with RH (*n=4*), Pru (*n=4*). **c)** Percentage of infection was calculated using HFFs grown on coverslips and stimulated with 10 U/mL IFNγ for 24 h. Following stimulation, HFFs were infected with either RH (MOI=1) or Pru (MOI=3) for another 24 h. Coverslips were fixed and stained with GRA7 for parasite PVM and Hoechst 33258 for nuclei. The number of PVs in 5-6 fields of the coverslips were counted for each condition. Experiments were performed with RH (*n=3*), Pru (*n=3*). **d)** Parasites per vacuole were determined 24 h p.i. with similar conditions and staining as described in (c) except MOIs used were 0.5 and 1 for RH (*n=3*) and Pru (*n=3*), respectively. **e)** Plaque assays were performed similarly as described for (a) with Pru WT (*n=20* for plaque number and *n=7* for plaque area), Pru∆*gra15* (*n=20* for plaque number and *n=7* for plaque area), Pru∆*gra15*+*GRA15* complemented (*n=12* for plaque number and *n=7* for plaque area). **f)** Plaque assays were performed similarly as described for (a) with RH (*n=5* for plaque number and *n=3* for plaque area) and RH+GRA15_||_ (*n=5* for plaque number and *n=3* for plaque area). **g)** Relative parasite growth was measured as described in (b) with Pru WT (*n=12*), Pru∆*gra15* (*n=12*), Pru∆*gra15*+*GRA15* complemented (*n=8*). **h)** Percentage of infection was calculated as described in (c) with Pru WT (*n=3*), Pru∆*gra15* (*n=3*), RH (*n=3*) and RH+GRA15_||_ (*n=3*). Statistical analysis was done by Two-way ANOVA followed by Tukey’s multiple comparison test. Data are represented as mean ± standard error of mean (SEM).

After invasion, *Toxoplasma* resides within the host cytosol in a PV and starts secreting GRAs into the PV lumen where they stay or get transported to the PVM or beyond the PVM into the host cell (Hakimi *et al*, 2017). The transport of GRAs beyond the PVM, but not onto the PVM, is mediated by a putative translocon containing the proteins MYR1/2/3 (Franco *et al*, 2016; Naor *et al*, 2018). However, Pru and Pru∆*myr1* parasites showed similar IFNγ-mediated reductions in plaque number and area (**Suppl. Fig. 1a**) indicating that these phenotypes are not influenced by GRAs secreted beyond the PVM. We therefore hypothesized that maybe a GRA in the PVM facing the host cytosol might be involved. GRA15 is a polymorphic *Toxoplasma* effector protein present in the PVM that activates the NF-κB transcription factor (Rosowski *et al*, 2011), a master regulator of cell signaling and cell death (Dutta *et al*, 2006), independent of MYR1 (Franco *et al*, 2016). The type I RH strain has an early stop codon in *GRA15* leading to a non-functional GRA15 (Rosowski *et al*, 2011). To determine if GRA15 plays a role in the susceptibility of Pru to IFNγ-mediated growth inhibition, we infected IFNγ-stimulated or naïve HFFs with Pru, Pru∆*gra15*, and the Pru∆*gra15* strain complemented with an HA-tagged copy of GRA15 (Pru∆*gra15*+*GRA15*) and measured plaque number and area after 6 days. Pru∆*gra15* parasites showed significantly less plaque number and area loss compared to wild-type parasites in IFNγ-stimulated HFFs (**Fig. 1e**). Complementation of ∆*gra*15 parasites with *GRA15* restored growth inhibition to wild-type levels (**Fig. 1e**). RH parasites expressing type II GRA15 (RH+GRA15_||_) (Rosowski *et al*, 2011) showed significantly more plaque loss compared to the wild-type RH strain (**Fig. 1f**). Furthermore, we observed that the enhanced GRA15-mediated susceptibility of Pru to IFNγ was already apparent 24 h post infection (p.i.) (**Fig. 1g**) and GRA15 also enhanced the IFNγ-mediated elimination of vacuoles (**Fig. 1h**).

It was recently shown that in immortalized HFFs, IFNγ-induced IDO1 expression determines the IFNγ-mediated growth inhibition of *Toxoplasma*. The MYR1-dependent secreted *Toxoplasma* effector TgIST was shown to protect *Toxoplasma* from IDO1-mediated growth inhibition in cells stimulated after infection by inhibiting STAT1-mediated *IDO1* expression but not in cells pre-stimulated with IFNγ (Bando *et al*, 2018). In contrast, we previously showed that IDO-mediated L-Trp degradation only plays a minor role in inhibition of parasite growth in IFNγ-stimulated primary HFFs (Niedelman *et al*, 2013). To rule out the role of IDO in the increased susceptibility of Pru parasites to IFNγ we show that L-Trp supplementation did not restore the reduction of plaque loss in RH and Pru strains (**Suppl. Fig. 1b**). Furthermore, there is no difference in IDO activity in RH vs. Pru-infected IFNγ-stimulated HFFs (**Suppl. Fig. 1c**). Thus, in IFNγ-stimulated HFFs parasite expression of GRA15 leads to reduced parasite growth and enhanced disappearance of vacuoles.

### GRA15 enhances IFNγ-induced endo-lysosomal fusion with the vacuole

In HUVEC cells, ubiquitin, ubiquitin-like proteins (LC3/GABARAP), and ubiquitin receptors (p62/NDP52) are recruited to the *Toxoplasma* PVM, eventually leading to its destruction by fusion with endo-lysosomes (Clough *et al*, 2016). To determine if this happens in HFFs and the potential role of GRA15 in this process, we pre-stimulated HFFs with IFNγ and infected cells with RH, Pru or Pru∆*gra15* parasites and measured accumulation of ubiquitin, p62, NDP52, LC3B, GABARAP and LAMP1 around the PVM 3 h p.i. Surprisingly, and unlike what has been observed in HeLa, HUVEC and murine cells (Clough *et al*, 2016; Haldar *et al*, 2015; Lee *et al*, 2015; Selleck *et al*, 2015), we observed that although PVMs of both RH and Pru strains were coated with ubiquitin in IFNγ-stimulated HFFs, a larger fraction of RH vacuoles was coated (**Fig. 2a**). We did not observe any difference in the ubiquitin coating intensity among the different parasite strains (**Suppl. Fig. 2a**). Deletion of *GRA15* had no effect on ubiquitination of Pru vacuoles (**Fig. 2a**). The type of ubiquitin linkage recruited to the PVM can influence the subsequent outcome (Swatek & Komander, 2016). We observed K63-linked, and no K48-linked (not shown), ubiquitin localized to the PVM (**Fig. 2b**). We observed a significantly larger fraction of Pru PVMs coated with p62 compared to RH (**Fig. 2c**). Deletion of *GRA15* from Pru resulted in significant fewer PVMs coated with p62 while a similar fraction of PVMs of the *GRA15* complemented strains as wild-type Pru were coated with p62 (**Fig. 2c**). Unlike the type I RH strain, the type I GT1 strain contains a functional GRA15, which we previously showed determines RH *vs.* GT1 differences in activation of NF-κB (Yang *et al*, 2013). Consistent with a role for GRA15 in mediating p62 PVM-recruitment, a significantly larger fraction of the vacuoles from the RH+GRA15_||_ and the GT1 strain stained positive for p62 compared to RH vacuoles (**Suppl. Fig. 2b**). Additionally, we observed that GT1 was significantly more susceptible to IFNγ-mediated parasite elimination compared to RH (**Suppl. Fig. 2c**). A similar fraction of PVMs of RH and Pru vacuoles contained NDP52 in IFNγ-stimulated HFFs (**Fig. 2d**). However, deletion of *GRA15* in Pru resulted in reduction of NDP52 recruitment to the PVM (**Fig. 2d**). Like p62, both LC3B and GABARAP were recruited to a larger fraction of the PVs of Pru compared to RH (**Fig. 2e and f**) in IFNγ-stimulated HFFs. The Pru∆*gra*15 strain had ~2-fold less vacuoles that were coated with LC3B and GABARAP compared to wild-type Pru (**Fig. 2e and f**). The *GRA15* complemented strain had a larger fraction of PVs coated with LC3B compared to the GRA15 deleted strain (**Fig. 2e**). To determine if the recruitment of LC3B, GABARAP and p62 is associated with lysosomal destruction of the vacuole we infected IFNγ-stimulated HFFs and counted LAMP1-positive vacuoles 3 h p.i. IFNγ enhanced the recruitment of LAMP1 to vacuoles of all strains but significantly more LAMP1-positive vacuoles were seen in Pru-infected, compared to RH-infected, cells. Deletion of *GRA15* significantly reduced the number of LAMP1-positive vacuoles (**Fig. 2g**). In many LAMP1-positive vacuoles the parasites were distorted, and they often did not stain positive for GRA7, used as parasite PV marker. However, by using the DNA-binding dye Hoechst these vacuoles still clearly contained parasite DNA but were in advanced stages of parasite degradation (**Suppl. Fig. 3**). The lysosomal inhibitor BafA1 significantly inhibited the disappearance of vacuoles in IFNγ-stimulated cells (**Fig. 2h**).

**Figure 2:**
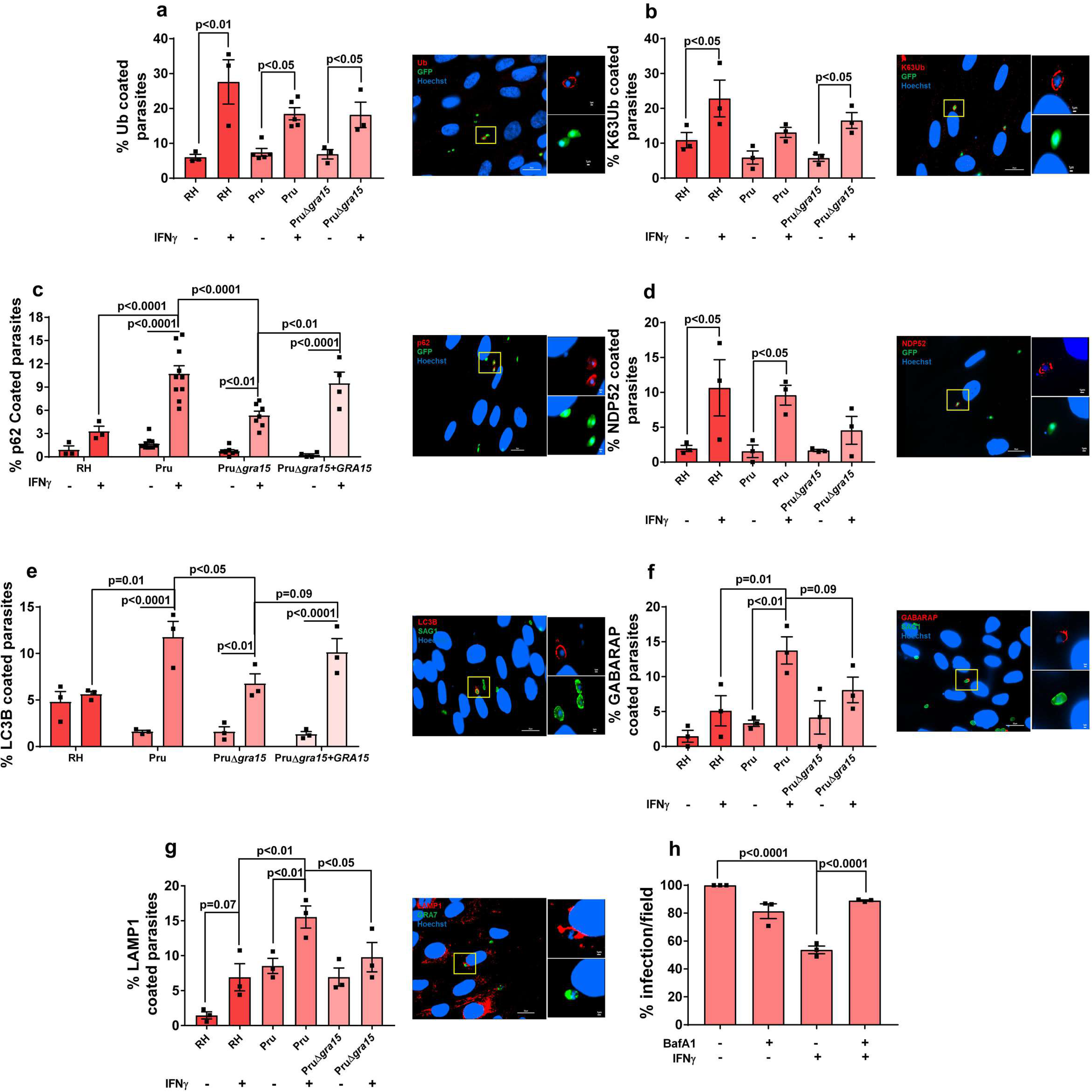
GRA15 enhances IFNγ-induced PVM decoration with autophagy-related proteins and endo-lysosomal-mediated vacuole destruction in HFFs. **a-g**) HFFs were stimulated with for 24 h with 10U/mL IFNγ or left unstimulated and subsequently infected with RH, Pru or Pru∆*gra*15 parasites for 3 h. The percentage of vacuoles that stained positive for **a)** Total ubiquitin (*n=3* for RH, *n=5* for Pru and *n=3* for Pru∆*gra15*), **b)** K63-linked ubiquitin (*n=3* for RH, *n=3* for Pru and *n=3* for Pru∆*gra15*), **c)** p62 (*n=3* for RH, *n=10* for Pru and *n=7* for Pru∆*gra15*, *n=4* for Pru ∆*gra15*+*GRA15*), **d)** NDP52 (*n=3* for RH, *n=3* for Pru and *n=3* for Pru∆*gra15*), **e)** LC3B (*n=3* for RH, *n=3* for Pru and *n=3* for Pru∆*gra15*, *n=3* for Pru ∆*gra15*+*GRA15*), **f)** GABARAP (*n=3* for RH, *n=3* for Pru and *n=3* for Pru∆*gra15*) and **g**) LAMP1 (*n=3* for RH, *n=3* for Pru and *n=3* for Pru∆*gra15*) is shown in the left bar diagram. On the right-hand side, a representative fluorescent image is shown for the *Toxoplasma* Pru strain, which expresses GFP. DNA was stained with Hoechst 33258. Scale bar is 10 μm. The yellow box inside each representative image is shown as inset pictures with magnification. **h)** The number of parasite-infected HFFs per 20X objective field was counted and compared between IFNγ-stimulated and IFNγ + bafilomycin A1 (100 nM) treated HFFs 24 h p.i. with Pru strain. Images from at least 6 fields were taken for each condition (*n=3*). Each dot represents one experiment. Each time at least 100 different vacuoles were scored and analyzed. Statistical analysis was done by Two-Way ANOVA followed with Tukey’s multiple comparison test (**a-g**) and One-way ANOVA for h. Data are represented as mean ± SEM.

Overall these results indicate that GRA15 enhances vacuole destruction via endo-lysosomal fusion in IFNγ-stimulated HFFs.

### GRA15-mediated enhancement of destruction of Pru vacuoles through endo-lysosomal fusion in IFNγ-stimulated HFFs correlates with PVM recruitment of p62, LC3B, GABARAP, and LAMP1 but not ubiquitin

In IFNγ-stimulated HUVEC cells only type II strain PVMs are ubiquitinated and this ubiquitination is indispensable for subsequent endo-lysosomal fusion and parasite elimination (Clough *et al*, 2016). However, in HFFs, strain differences in ubiquitination of the PVM (**Fig. 2a, 2b**) did not correlate with recruitment of p62, LC3B, GABARAP and LAMP1 to the PVM (**Fig. 2c-g**). Furthermore, when we inhibited ubiquitination using a specific inhibitor of E1 ubiquitin activating enzymes (PYR 41), we observed a significant reduction in the fraction of PVMs coated with ubiquitin but no effect on the fraction of PVMs coated with p62 or LC3B (**Fig. 3a**). Thus, the fraction of PVMs containing p62 or LC3B does not correlate with the fraction of PVMs containing ubiquitin. Consistent with this, only 25% of the vacuoles were coated with both ubiquitin and p62, only 40% of the vacuoles were coated with both ubiquitin and LC3B and only 20% of the vacuoles were coated with ubiquitin and GABARAP (**Fig. 3b-d**). In contrast, 90% of the vacuoles were coated with p62 and LC3B (**Fig. 3e**) and 78% with p62 and GABARAP (**Fig. 3f**). Given that ~20% of the vacuoles that are coated with only p62 and ~60% are only Ub positive (**Fig. 3b**) these data suggest that recruitment of p62 to the PVM is not dependent on PVM ubiquitination.

**Figure 3:**
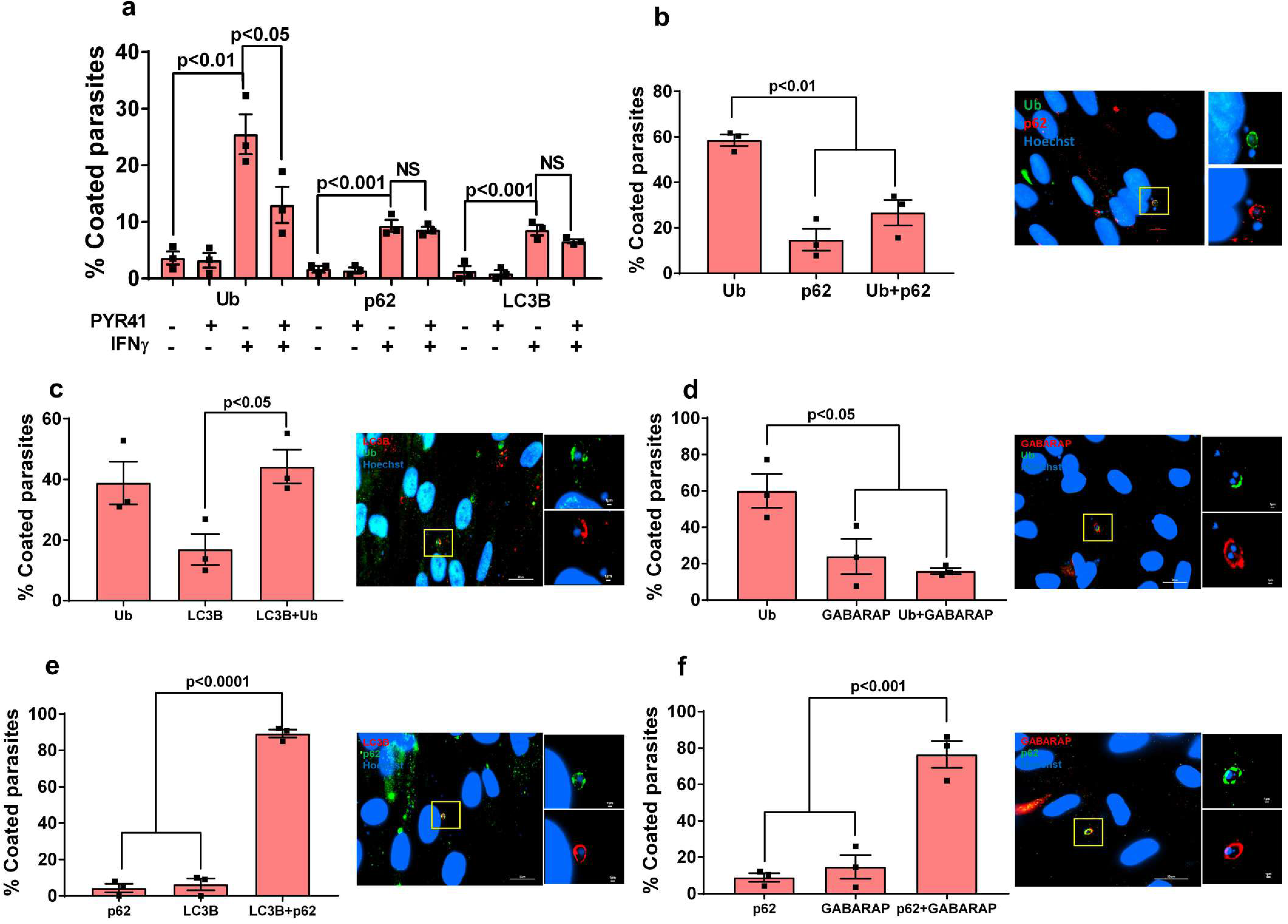
PVM decoration with autophagy adaptors correlates with p62 but not ubiquitination. **a)** HFFs were stimulated with IFNγ (10 U/mL for 24 h) or left unstimulated and subsequently treated with PYR41 (1 μM) for 2 h prior to infection. Cells were washed and subsequently infected with Pru parasites (MOI=3) for 3 h. The percentage of vacuoles that stained positive for total ubiquitin, p62 or LC3B was determined (*n=3*). **b-f**) For all the co-staining experiments IFNγ-stimulated HFFs infected for 3 h with the Pru strain were used. For each staining at least 50 vacuoles were scored (*n=3*). On the right-hand side, a representative fluorescent image is shown for the *Toxoplasma* Pru strain. Scale bar is 10 μm. The yellow box inside each representative image is shown as inset pictures with higher magnification. The total number of coated vacuoles was set at 100% and the percentage of vacuoles positive for Ub, p62 and/or LC3B/GABARAP was calculated. Statistical analysis was done by One-way ANOVA followed with Tukey’s multiple comparison test. Data are represented as mean ± SEM.

### GRA15 binds TRAFs and recruits TRAF6 and ubiquitin-like molecules and receptors to the parasitophorous vacuole to mediate parasite elimination

We wanted to determine if GRA15 inhibits parasite growth in IFNγ-stimulated HFFs through its ability to activate the NF-κB transcription factor (Rosowski *et al*, 2011). To test this, we treated HFFs with BAY11-7082, a known inhibitor of NF-κB activation (García *et al*, 2005), 2 h before infection and 24 h p.i. measured parasite growth. Addition of BAY11-7082 (1 μM) did not restore parasite growth in IFNγ-stimulated HFFs (**Fig. 4a**). At this concentration, BAY11-7082 can successfully inhibit the activation of NF-κB triggered by Pru parasites (**Fig. 4b)**. BAY11-7082 was reported to inhibit NF-κB activation via inhibition of E2 ubiquitin conjugating enzymes that mediate K63- and linear polyubiquitin chains (Strickson *et al*, 2013). We therefore also counted the percentage of PVs coated with ubiquitin or p62 in IFNγ-stimulated HFFs and observed that although treatment with BAY11-7082 decreased the fraction of PVs that were coated with ubiquitin, it had no effect on the fraction of PVs coated with p62 (**Suppl. Fig. 4a**).

**Figure 4:**
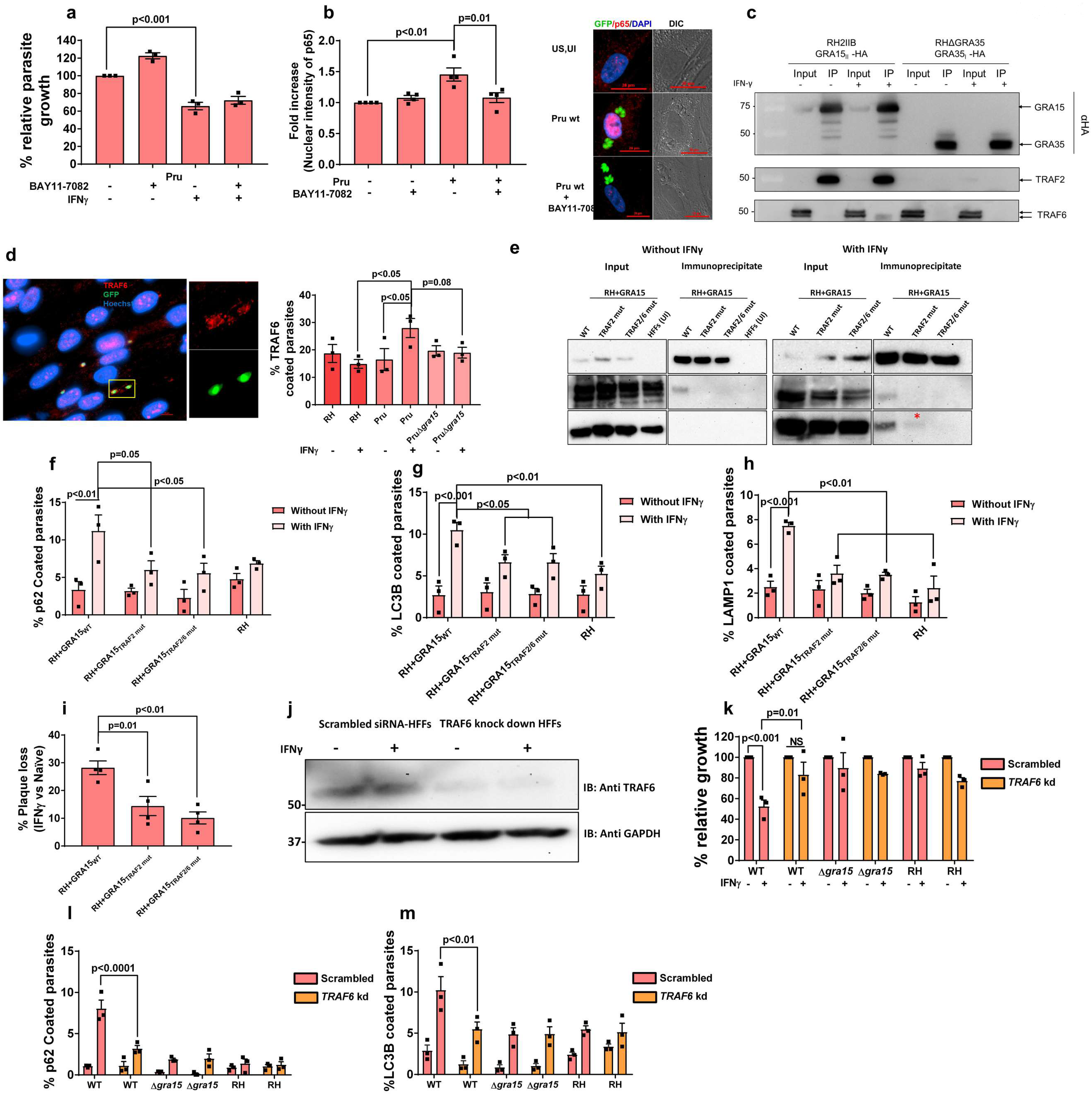
GRA15-mediated parasite growth reduction in IFNγ-stimulated HFFs is independent of its ability to activate NF-κB but dependent on GRA15’s interaction with the E3 ubiquitin ligases TRAF2 and TRAF6. **a)** HFFs were stimulated with IFNγ for 24 h (10 U/ml) or left unstimulated. The NF-κB inhibitor BAY11-7082 (1μM) was added 2 h pre-infection and HFFs were subsequently infected with Pru parasites. Parasite growth (using luciferase assay) was measured 24 h p.i. Means from unstimulated cells were set at 100%. Experiments were performed three times. **b**) Nuclear translocation of the NF-κB p65 subunit was quantified in HFFs or HFFs treated with BAY11-7082 (1μM, added 2 h pre-infection) 24 h p.i. with Pru parasites. Experiments were done 4 times where each dot represents one experimental mean of at least 15 nuclei. In the right panel, representative images are shown. Parasites were expressing GFP, nuclei are stained with Hoechst 33258. Scale bar is 20 μm. **c)** Immunoprecipitation and Western blot were performed in HFFs with and without IFNγ (10 U/ml) using an RH strain expressing type II GRA15-HA and as a control RH expressing GRA35-HA. The blots using antibodies against TRAF2 and TRAF6 were made after stripping the first blot. The inputs loaded represent 1% of total lysate prepared for immunoblotting and mass spec (Suppl. Table 2). The antibodies against TRAF2 and TRAF6 were obtained from Santa Cruz Biotechnology (Suppl. Table 1). Full length blots for this figure can be observed in supplementary figures. 4c-e. **d)** Immunofluorescence analysis of TRAF6 recruitment to the PVM of HFFs infected for 3 h with RH, Pru and Pru∆*gra*15 strains. On the right-hand side, a representative fluorescent image is shown of TRAF6 recruitment to the PVM where *Toxoplasma* Pru strain expresses GFP, DNA was stained with Hoechst 33258. Scale bar is 10 μm. *n=3* for all the strains. The antibody against TRAF6 was purchased from Abnova (Suppl. Table 1). **e)** Immunoprecipitation and Western blot were performed in HFFs with and without IFNγ (10 U/ml) using a RH+GRA15_WT_, RH+GRA15_TRAF2mut_ or RH+GRA15_TRAF2/6mut_. For the unstimulated cells, uninfected HFFs were used an additional negative control (left panel). Left panel and right panel were run on a single gel; vertical white lines indicate excision of irrelevant lanes. Full length blots are in supplementary figures 4f-k. The antibodies used against TRAF2 and TRAF6 were purchased from Cell Signaling Technology and Abnova, respectively. The asterisks (*) in the lower right panel indicates the faint band of TRAF6 in the RH+GRA15_TRAF2mut_ immunoprecipitate. **f-h)** Immunofluorescence analysis of p62, LC3B and LAMP1 with and without IFNγ in RH, RH+GRA15_WT_. All the experiments were done 3 times with each of the strains. **i)** Plaque assays were performed with RH+GRA15_WT_, RH+GRA15_TRAF2mut_ or RH+GRA15_TRAF2/6mut_ (*n=3*). **j)** Expression of TRAF6 was detected by Western blotting of lysates from scrambled siRNA transfected and TRAF6-specific siRNA transfected HFFs with and without IFNγ. **k)** Relative parasite growth was measured in scrambled siRNA transfected HFFs and *TRAF6* knockdown HFFs using luciferase based assay with indicated strains with and without IFNγ (*n=3*). The antibody against TRAF6 used here was from Abcam (Suppl. Table 1). Full length blots are in supplementary figures 4l-m. **l-m)** Immunofluorescence analysis of p62 and LC3B was done in scrambled siRNA transfected HFFs and *TRAF6* knockdown HFFs with and without IFNγ using indicated strains (*n=3*). Statistical analysis was done by One-way ANOVA followed with Tukey’s multiple comparison test (a, b and i) and two-Way ANOVA followed with Tukey’s multiple comparison test (d, f-h and k-m). Data are represented as mean ± SEM.

To identify the mechanism by which GRA15 enhanced the recruitment of ubiquitin-like molecules and ubiquitin receptors to the PVM, we immunoprecipitated GRA15 from naive and IFNγ-stimulated HFFs 8 h p.i. As a control we immunoprecipitated GRA35, which we and others recently showed is another PVM-localized GRA (Nadipuram *et al*, 2016; Wang *et al*, 2019b). GRA15 specifically immunoprecipitated multiple ubiquitin ligases, TRAF1, TRAF2, BIRC2, BIRC3, and TNFAIP3 (also named A20), while in IFNγ-stimulated cells also polyubiquitin and TRAF6 were immunoprecipitated (**Suppl. Table 2**). To confirm some of these results, we immunoprecipitated GRA15 and blotted for TRAF2 and TRAF6. Indeed GRA15, but not GRA35, immunoprecipitated TRAF2 in both stimulated and unstimulated HFFs while a small amount of TRAF6 was only immunoprecipitated in IFNγ-stimulated HFFs (**Fig. 4c)**. We also observed a significantly larger fraction of Pru PVs coated with TRAF6 compared to RH PVs in IFNγ-stimulated HFFs (**Fig. 4d**). Pru∆*gra15* PVMs had less recruitment of TRAF6 compared to the wild-type Pru strain (**Fig. 4d**). However, this antibody seemed to have quite some background staining. TRAF6 is a ring type E3 ubiquitin ligase and catalyzes the formation of K63-linked ubiquitination (Deng *et al*, 2000) and was reported to be associated with ubiquitination of PVs in murine cells (Haldar *et al*, 2015). We observed that although inhibition of TRAF6 ubiquitin ligase activity significantly lowered K63-linked ubiquitination on the PVM this had no effect on recruitment of p62 to the PVM (**Suppl. Fig. 4b)**. Because inhibiting ubiquitin activating (E1), ubiquitin conjugating (E2) and ubiquitin ligase (E3) activity did not affect p62 recruitment, if TRAF6 has a role in vacuole destruction it is unlikely to be the ubiquitination of the PVM but more likely the recruitment of p62 via its p62 binding domain. Recently, we have reported that activation of NF-κB by GRA15 depends on in its interaction with TRAF2 as well as TRAF6 in HEK cells (Sangaré *et al*, 2019). GRA15 has 2 binding motifs for TRAF2 and one for TRAF6 (**Suppl. Fig. 5a**) and contains at least 4 high confidence ubiquitination sites (**Suppl. Fig. 5b**). However, none of the ubiquitination sites have the consensus downstream p62-TRAF6 binding sites (**Suppl. Fig. 5c**). To directly determine the relevance of the GRA15 interaction with TRAF2 and TRAF6 at the PVM, we generated RH parasites that expressed either HA-tagged wild-type GRA15 (RH+GRA15_WT_) or GRA15 with mutated TRAF2-(RH+GRA15_TRAF2mut_) or TRAF2- and TRAF6-binding sites (RH+GRA15_TRAF2/6mut_). Immunofluorescence and Western blot analysis showed that GRA15 had a similar PV/PVM localization in RH+GRA15_WT_, RH+GRA15_TRAF2mut_, and RH+GRA15_TRAF2/6mut_ and had similar expression levels in these parasites (**Suppl. Fig. 5e and f**). We immunoprecipitated GRA15 from these different parasites from lysates generated from naive or IFNγ-stimulated infected HFFs 8 h p. i. Immunoprecipitated GRA15_WT_ pulled down TRAF2 from infected HFF lysates while no TRAF2 was pulled down after immunoprecipitation of GRA15_TRAF2mut_ or RH+GRA15_TRAF2/6mut_. In the lysate from IFNγ-stimulated infected HFFs, GRA15_WT_ pulled down TRAF2 and TRAF6 but no TRAF2 or TRAF6 was pulled down after immunoprecipitation of GRA15_TRAF2/6mut_ and no TRAF2 and significantly less TRAF6 was pulled down after immunoprecipitation of GRA15_TRAF2mut_. These data confirmed that mutation of the respective TRAF2/6-binding sites indeed abrogated the binding to TRAF2 or to TRAF2 and TRAF6 (**Fig. 4e, Suppl. Fig. 5d**). The fact that less TRAF6 was immunoprecipitated with the GRA15_TRAF2mut_ compared to GRA15_WT_ could indicate that TRAF6 was coming down via binding to TRAF2(Davies *et al*, 2005). We observed that significantly more PVs of RH+GRA15_WT_ contained p62 (**Fig. 4f**), LC3B (**Fig. 4g**) and LAMP1 (**Fig. 4h**) compared to the PVs of RH+GRA15_TRAF2mut_ parasites. Recruitment of p62, LC3B and LAMP1 to the PVM of RH+GRA15_TRAF2mut_ and RH+GRA15_TRAF2/6mut_ parasites was similar, indicating that this recruitment was mainly mediated via the GRA15 TRAF2-binding sites (**Fig. 4f-h**). Furthermore, IFNγ-mediated killing of RH+GRA15_WT_ was significantly higher compared to RH+GRA15_TRAF2mut_ or RH+GRA15_TRAF2/6mut_ parasites (**Fig. 4i**).

Because TRAF2 does not contain a p62-binding domain we hypothesized that its main role was the recruitment of TRAF6. To determine the role of TRAF6, we knocked down TRAF6 in HFFs, using a stable lentivirus mediated siRNA system (**Fig. 4j**). The growth of Pru parasites in presence of IFNγ in these TRAF6 knockdown HFFs was significantly restored compared to scrambled siRNA transfected HFFs (**Fig. 4k**) whereas both RH and Pru∆*gra15* were resistant in both cell types (**Fig. 4k**). Furthermore, recruitment of p62 and LC3B in IFNγ-stimulated cells was significantly less in TRAF6 knockdown HFFs compared to scrambled siRNA transfected HFFs (**Fig. 4l and m**). These results indicate that GRA15 by binding with ubiquitin ligases TRAF2 and TRAF6 mediates the susceptibility of type II Pru strains in IFNγ-stimulated primary human fibroblasts.

### GRA15 mediates susceptibility of type II parasites to IFNγ-mediated killing in murine fibroblasts by binding to TRAF6

Our results show that GRA15 mediates Pru susceptibility to IFNγ in HFFs by enhancing lysosomal destruction of the vacuole. In IFNγ-stimulated MEFs, type II GRA15 plays a role in recruitment of GBPs to the PVM via an unknown mechanism (Fisch *et al*, 2019; Virreira Winter *et al*, 2011). In IFNγ-stimulated MEFs, GBP recruitment is initiated by PVM recruitment of so-called ‘pioneer’ IRGs such as IRGB6 (Haldar *et al*, 2015). In MEFs recruitment of TRAF6 and TRIM21, two ubiquitin ligases, to the PVM enhances subsequent recruitment of IRGs and GBPs that target the vacuole for degradation (Haldar *et al*, 2015; Foltz *et al*, 2017). We hypothesized that TRAF6 recruitment by GRA15 might mediate the recruitment of GBPs and IRGs as TRAF6 has a p62 binding motif and p62 has an LC3 interacting region (LIR) motif that can mediate recruitment of LC3B and GABARAP (Lamark *et al*, 2017). To determine the role of GRA15 in MEFs, we infected IFNγ-stimulated MEFs and measured PV recruitment of TRAF6, IRGB6, ubiquitin (K63 and K48), GBPs, and LC3B. As demonstrated by others (Foltz *et al*, 2017; Haldar *et al*, 2015), Pru parasites had a significantly larger fraction of its vacuoles coated with these markers compared to RH parasites. This enhanced PVM recruitment was partially mediated by GRA15 as we observed significantly fewer PVs coated with these markers in Pru∆*gra15* parasites (**Fig. 5a-f**). In contrast to what we observed in HFFs, significantly more Pru vacuoles were coated with ubiquitin, predominantly K63-linked, in IFNγ-stimulated MEFs compared to RH vacuoles which is in agreement with previous studies (Foltz *et al*, 2017; Haldar *et al*, 2015). We determined parasite growth in IFNγ-stimulated MEFs 24 h p.i. and observed that Pru∆*gra15* had significantly less IFNγ-mediated growth reduction compared to either wild type or GRA15 complemented parasites (**Fig. 5g**). Pru∆*gra15* also showed significantly less plaque loss compared to its parental or complemented strains (**Fig. 5h**). RH expressing type II GRA15 formed less and smaller plaques compared to wild-type RH in IFNγ-stimulated MEFs (**Fig. 5i-j**).

**Figure 5:**
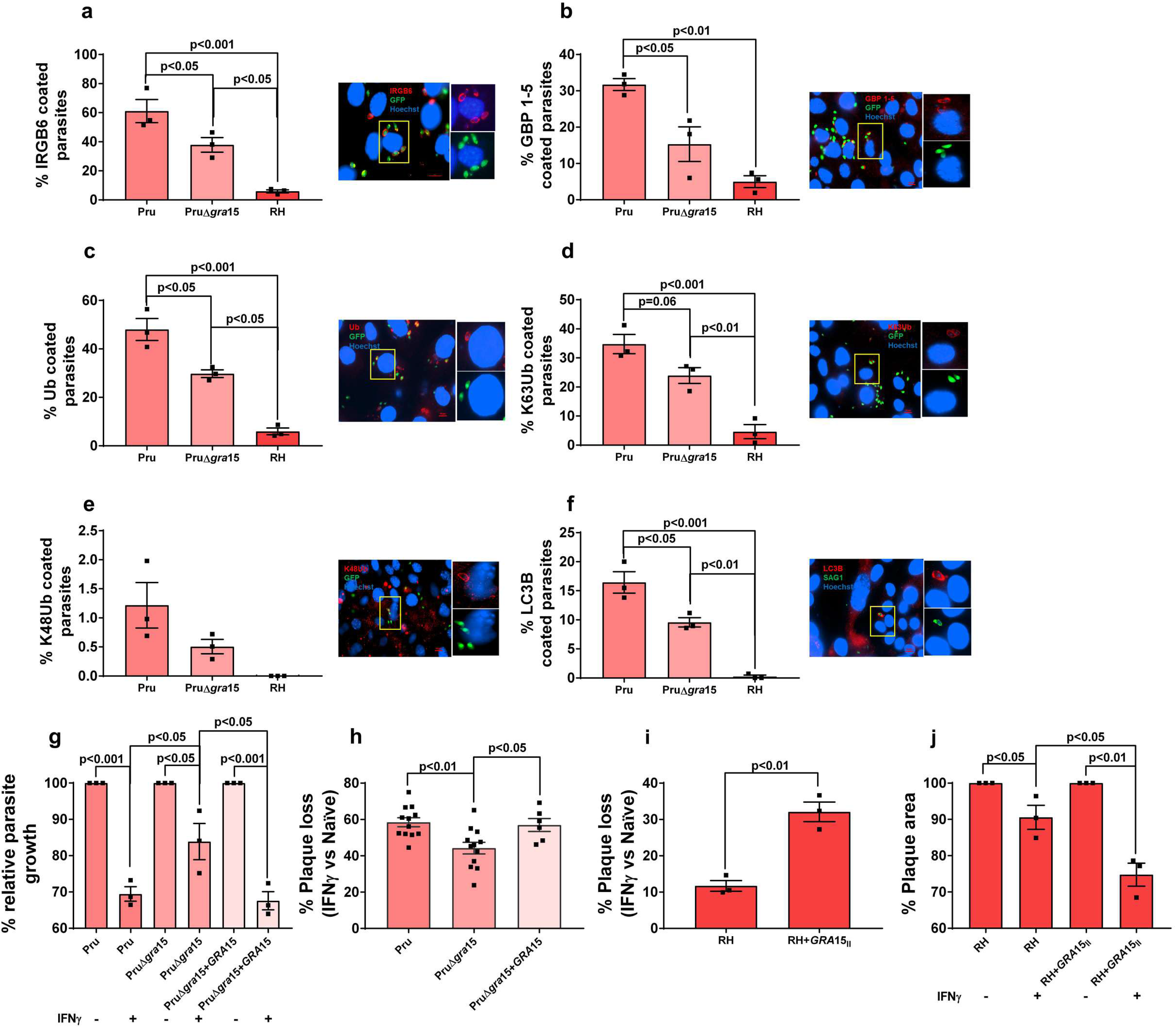
GRA15 enhanced susceptibility of type II strains to IFNγ-mediated killing in MEFs correlates with enhanced recruitment of IRGB6, GBPs, Ubiquitin, and LC3B. MEFs were stimulated with IFNγ for 24 h (100 U/ml) **a-f)** IFNγ-stimulated MEFs were infected with Pru, Pru∆*gra*15 or RH for 3 h and subsequently fixed, permeabilized and stained for **a)** IRGB6, **b)** GBPs, **c)** total ubiquitin, **d)** K63-linked ubiquitin, **e)** K48-linked ubiquitin and **f)** LC3B. For analysis, at least 100 vacuoles were scored. All experiments were performed 3 times. On the right-hand side, a representative fluorescent image is shown for the *Toxoplasma* Pru strain, which expresses GFP. DNA was stained with Hoechst 33258. Scale bar is 10 μm. The yellow box inside each representative image is shown as an inset picture with magnification. **g)** MEFs were stimulated with IFNγ for 24 h (100 U/ml) or left unstimulated and subsequently infected with Pru, Pru∆*gra*15 and *GRA*15 complemented parasites for 24 h for measuring parasite growth by luciferase assay. All the experiments from **a-g** were performed 3 independent times. **h)** Plaque numbers were measured for Pru (*n=12*), Pru∆*gra*15 (*n=12*) and *GRA15* complemented parasites (*n=6*). **I-j)** MEFs were stimulated with IFNγ for 24 h (100 U/ml) or left unstimulated and subsequently infected with RH or RH expressing type II GRA15. Plaque numbers were counted (*n=3*), and areas were measured 4 days p.i. (*n=3*). Statistical analysis was done by One-way ANOVA followed with Tukey’s multiple comparison test (a-h), except for (g, j) for which Two-way ANOVA followed with Tukey’s multiple comparison test was performed whereas for (i) two sample student’s t test was performed. Data are represented as mean ± SEM.

To establish that the GRA15-enhanced recruitment of these markers to the vacuole could be mediated by its interaction with TRAF6, we immunoprecipitated GRA15 from naïve and IFNγ-stimulated MEFs and performed Western blot for TRAF2 and TRAF6. We observed that akin to HFFs, GRA15 in MEFs also binds TRAF2 and TRAF6 (**Fig. 6a**). We compared recruitment of TRAF6 to PVs of RH, Pru and Pru∆*gra15* in MEFs. Akin to HFFs, we observed that in IFNγ-stimulated MEFs, significantly more Pru PVs contained TRAF6 (3-fold increase) compared to either RH or Pru∆*gra15* PVs (**Fig. 6b**). Significantly fewer Pru PVs were coated (3-fold decrease) with IRGB6 in in *Traf6*^−/−^ MEFs compared to wild-type MEFs (**Fig. 6c**). Furthermore, the difference in IRGB6 coating between Pru and Pru∆*gra15* PVMs in wild-type MEFs disappeared in *Traf6*^−/−^ MEFs (**Fig. 6c**). We also measured the growth of Pru, Pru∆*gra15* and *GRA15* complemented parasites in wild type, *Traf6*^−/−^, and NF-κB *p65*^−/−^ MEFs 24 h p.i. In wild type and NF-κB *p65*^−/−^ MEFs both Pru and GRA15 complemented parasites showed a significant growth reduction upon IFNγ-stimulation (**Fig. 6d**). In contrast, in *Traf6*^−/−^ MEFs, the action of IFNγ was abolished and GRA15-expressing parasites no longer showed reduced growth compared to the Pru∆*gra15* strain, which was resistant to IFNγ in all MEFs (**Fig. 6d**). To further establish the role of GRA15-mediated TRAF6 binding in parasite growth inhibition in IFNγ-stimulated MEFs we infected MEFs with RH, RH+GRA15_WT_, RH+GRA15_TRAF2mut_ or RH+GRA15_TRAF2/6mut_ and enumerated the vacuoles with recruitment of IRGB6, GBP1-5, p62, TRAF6 and ubiquitin. By immunoprecipitation of GRA15 and Western blotting for TRAF2 and TRAF6, we demonstrated that these mutants are indeed unable to bind TRAF2 or TRAF2 and TRAF6 (**Fig. 6e**). Furthermore, as expected, the RH+GRA15_TRAF2/6mut_ strain no longer recruited TRAF6 to the vacuole while TRAF6 recruitment to the PVM of RH+GRA15_TRAF2mut_ was similar to RH+GRA15_WT_ (**Fig. 6f**). The reduced recruitment of TRAF6 to the PVM of the RH+GRA15_TRAF2/6mut_ strain was associated with a reduced recruitment of IRGB6, GBP1-5 and p62 (**Figure 6g-i**). In contrast, mutating the TRAF2 or the TRAF2 and TRAF6 binding site on GRA15 did not affect ubiquitin coating of the PVM (**Fig. 6j**) suggesting that although GRA15 expression is associated with increased ubiquitin coating (**Fig. 5c, d**) this is not mediated by binding of TRAF6 or TRAF2 to GRA15. The RH+GRA15_TRAF2/6mut_ strain was also less susceptible to IFNγ compared to either the RH+GRA15_WT_, RH+GRA15_TRAF2mut_ strains as indicated by reduced plaque loss upon IFNγ-stimulation (**Fig. 6k**).

**Figure 6:**
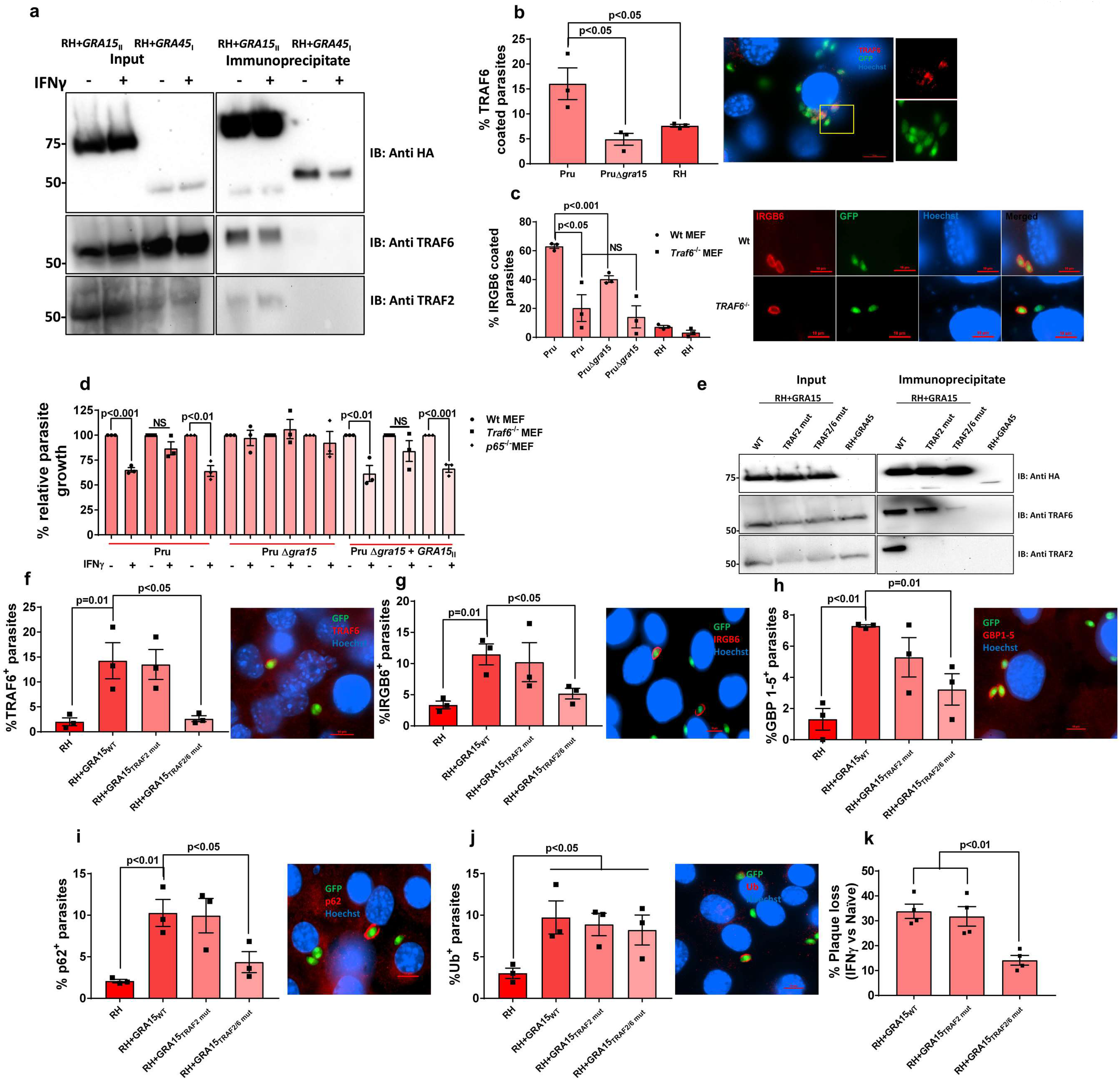
GRA15 enhanced susceptibility of the type II Pru strain in IFNγ-stimulated MEFs is dependent on TRAF6. **a)** Immunoprecipitations and Western blots were performed on MEFs stimulated or not with IFNγ (100 U/ml) and infected with an RH strain expressing type II GRA15-HA and as a control RH expressing GRA45-HA. The blots using antibodies against TRAF6 and TRAF2 were made after stripping the first blot. Left panel and right panel were run on a single gel; vertical white lines indicate excision of irrelevant lanes. Full length blots are in supplementary figures 6a-c. The antibodies used against TRAF2 and TRAF6 were purchased from Cell Signaling Technology and Abnova, respectively. **b)** IFNγ-stimulated MEFs were infected with Pru, Pru∆*gra*15 and RH for 3 h and subsequently stained for TRAF6. On the right-hand side, a representative fluorescent image is shown for the *Toxoplasma* Pru strain, which expresses GFP. DNA was stained with Hoechst 33258. Scale bar is 10 μm. The yellow box inside each representative image is shown as an inset picture with magnification. Experiments were performed 3 times. **c)** IRGB6 staining on PVM after infection of wild-type or *Traf6* ^−/−^ MEFs for 3 h with Pru, Pru∆*gra*15 and RH. On the right-hand side, a representative fluorescent image is shown of IRGB6 coating on the PVM of both wild-type and *Traf6* ^−/−^ MEF. At least 100 different vacuoles were observed and analyzed for each experiment (*n=3*). **d)** Parasite growth was measured 24 h p.i. using luciferase readout from unstimulated and IFNγ-stimulated wild-type, NFκB *p65*^−/−^ and *Traf6* ^−/−^ MEFs. Growth was compared between Pru, Pru∆*gra*15 and Pru∆*gra*15+*GRA*15 (complemented) strains. Reading from unstimulated cells was considered as 100% and percentage growth in IFNγ-stimulated cells was expressed relative to unstimulated cells. Experiments were performed three times with each MEF type. **e)** Immunoprecipitation and Western blot were performed in MEFs infected with RH+GRA15_WT_, RH+GRA15_TRAF2mut_ or RH+GRA15_TRAF2/6mut_ and as a control RH expressing GRA45-HA. Left panel and right panel were run on a single gel; vertical white lines indicate excision of irrelevant lanes. Full length blots are in supplementary figures 6d-f. The antibodies used against TRAF2 and TRAF6 were purchased from Cell Signaling Technology and Abnova, respectively. **f-j)** Immunofluorescence analysis of TRAF6, IRGB6, GBP1-5, p62 and ubiquitin was done in IFNγ-stimulated MEFs infected with RH+GRA15_WT_, RH+GRA15_TRAF2mut_ or RH+GRA15_TRAF2/6mut_ (*n=3*). **k)** Plaque assays were performed with RH+GRA15_WT_, RH+GRA15_TRAF2mut_ or RH+GRA15_TRAF2/6mut_ (*n=4*). Statistical analysis was done by two samples student’s t test (f-j) and One-way ANOVA followed with Tukey’s multiple comparison test for b and k and Two-way ANOVA for c and d. Data are represented as mean ± SEM.

Thus, GRA15, by recruiting TRAF6 in IFNγ-stimulated cells, and not by activating NF-κB p65, enhanced parasite susceptibility to IRG/GBP-dependent elimination in MEFs.

## Discussion

*Toxoplasma* strain-dependent susceptibility to IFNγ in different human and murine cells *in vitro* is well established (Bando *et al*, 2018; Clough *et al*, 2016; Haldar *et al*, 2015; Niedelman *et al*, 2012; Qin *et al*, 2017; Selleck *et al*, 2013, 2015; Niedelman *et al*, 2013). However, *Toxoplasma* effectors that affect strain differences in susceptibility to IFNγ-induced cell autonomous immunity in human cells have not been described. We show that in IFNγ-stimulated primary HFFs the *Toxoplasma* effector GRA15 (Jensen *et al*, 2011; Rosowski *et al*, 2011) mediates the recruitment of the E3 ubiquitin ligase TRAF6, its binding partner p62 and LC3B and GABARAPs, eventually leading to endo-lysosomal fusion with the vacuole and parasite elimination.

It was previously shown that in murine cells *Toxoplasma* GRA15 enhances the recruitment of GBP1-5 to the vacuole (Fisch *et al*, 2019; Virreira Winter *et al*, 2011). However, it was a mystery how GRA15 mediated this recruitment of GBP1-5 as its only known function was the activation of the NF-κB transcription factor, which does not take place until four hours after infection, while its effect on GBP recruitment can be seen within 1 hour (Rosowski *et al*, 2011; Virreira Winter *et al*, 2011). Here we solve this mystery by showing that in both HFFs and MEFs, GRA15 enhances the recruitment of TRAF6 to the PVM. TRAF6 was previously shown to be required for subsequent recruitment of p62, further ubiquitination of the vacuole and its eventual destruction by IRGs and GBPs (Haldar *et al*, 2015). We confirmed those data and additionally showed that in *Traf6*^−/−^ MEFs the difference in susceptibility between Pru and Pru∆*gra15* disappears further confirming that the effect of GRA15 is mediated through TRAF6. Thus, GRA15 enhances IFNγ-mediated parasite elimination in both HFFs and MEFs although the exact mechanism of vacuole elimination is different (**Fig. 7**).

**Figure 7:**
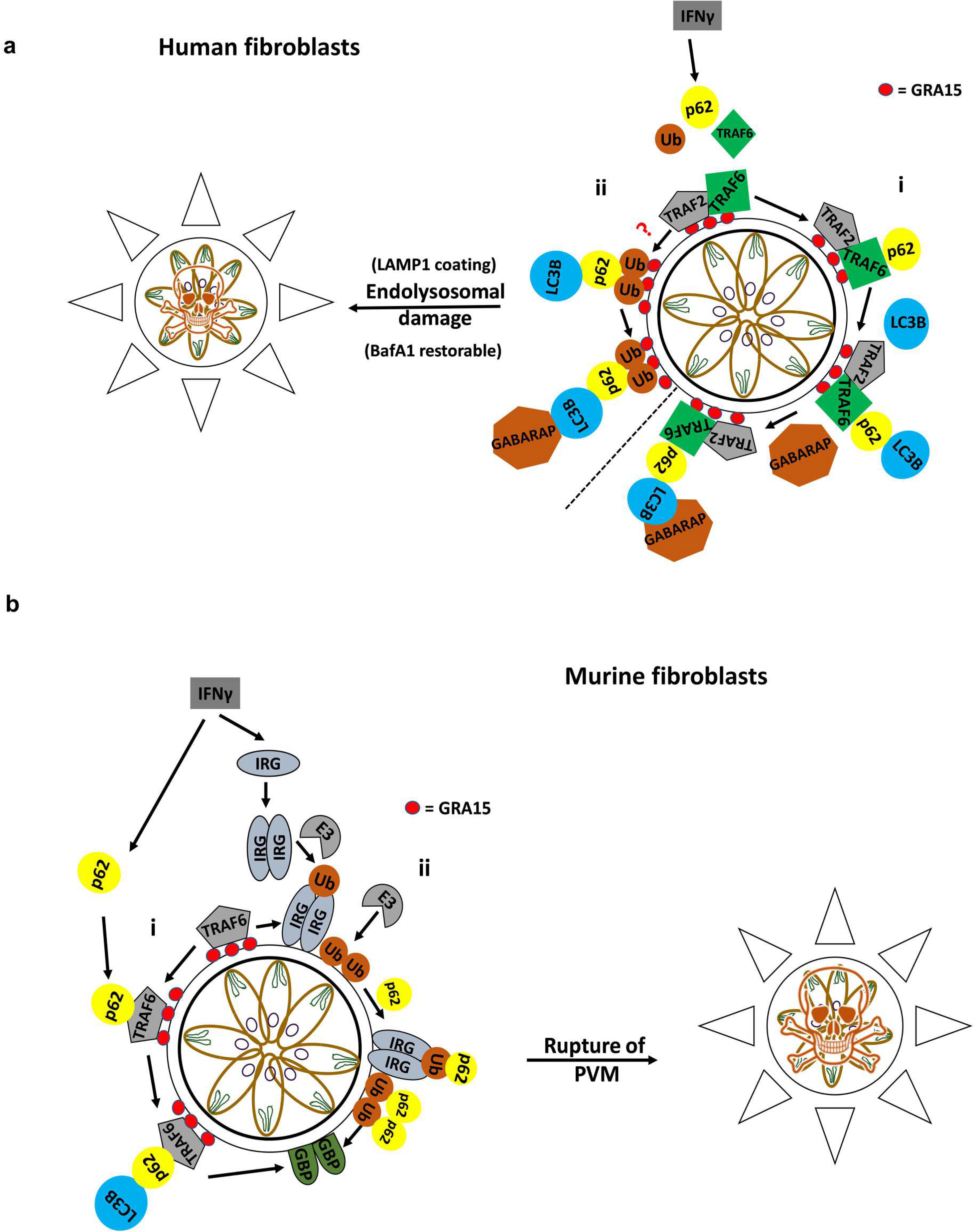
Proposed model of GRA15-mediated susceptibility to IFNγ-induced growth inhibition in human and murine fibroblasts. **a)** In HFFs, IFNγ-stimulation causes recruitment of TRAF2/6 to GRA15 on the PVM of *Toxoplasma*. What determines the IFNγ-induced binding of TRAF6 to GRA15 is unknown, although our results indicate TRAF2 plays a role in this. (i) Recruitment of TRAF6 mediates further recruitment of p62, LC3B and GABARAP. This subsequently leads to endo-lysosomal fusion with the PVM and elimination of the parasites. (ii) On the other hand, IFNγ-stimulation induced recruitment of ubiquitin to the PVM which could also recruit its adaptor proteins p62 and subsequently LC3B and GABARAP, which together can lead to endo-lysosomal fusion with the PVM and help in elimination of the parasites. **b)** In MEFs, GKS IRGs (IRGB6 in this study) are recruited to the PVM upon IFNγ-stimulation. We observed a decrease in IRG loading in *Traf6*^−/−^ MEFs suggesting that TRAF6 might act upstream of IRG loading. However, these IRGs also recruit other E3 ubiquitin ligases which can cause PVM ubiquitination which in turn could recruit p62. GRA15 by binding to TRAF6 accelerates the process and recruits more p62, LC3B and IRGs. This subsequently leads to enhanced recruitment of GBPs and damage to the PVM which ultimately kills the parasites.

Previously it was shown that ubiquitination of the PVM is a strictly strain-dependent phenomenon (Clough *et al*, 2016; Haldar *et al*, 2015; Lee *et al*, 2015; Selleck *et al*, 2015) where initial ubiquitination recruits p62 which further recruits the E3 ubiquitin ligases TRAF6 and TRIM21 to generate an amplification loop that recruits further p62 and LC3 eventually controlling parasite growth (Foltz *et al*, 2017; Haldar *et al*, 2015). However, our data show that in contrast to what has been observed in MEFs, PVM ubiquitination is not strain specific in HFFs, inhibition of PVM ubiquitination has no effect on p62 or LC3B recruitment, and a significant fraction of p62-coated PVMs do not contain ubiquitin. Also, in MEFs infected with RH or RH parasites expressing wild-type GRA15 or GRA15 TRAF-binding mutants the recruitment of ubiquitin to the PVM did not correlate with recruitment of TRAF6, P62, IRGB6 or GBPs. Our data suggest that the PVM-localized GRA15 parasite effector recruits TRAF6, which then likely further recruits adaptor proteins that in other cell types appear to be recruited by PVM ubiquitination (Clough *et al*, 2016; Selleck *et al*, 2015). In HeLa cells the recruitment of p62 and other markers caused parasite growth stunting but no vacuole destruction (Selleck *et al*, 2015) whereas in HUVEC cells, vacuole destruction by endo-lysosomal fusion was described to be ubiquitin and p62 dependent (Clough *et al*, 2016). In HFFs we showed that p62 is recruited to the PVM, possibly via its TRAF6-binding domain, and parasite vacuole destruction occurs through endo-lysosomal fusion. What determines the differences between these different cell types is currently unknown.

Although in MEFs GRA15 significantly increased the percentage of vacuoles targeted by IRGB6, vacuoles of the Pru∆*gra15* strain were still significantly more targeted compared to the RH strain. This is likely due to strain differences in *ROP5,* as we and others previously showed that II strains have *ROP5* alleles that are less effective at inhibiting IRG loading onto the PVM and that expression of type I or type III *ROP5* alleles significantly reduce coating of type II vacuoles with IRGB6 (Etheridge *et al*, 2014; Fleckenstein *et al*, 2012; Niedelman *et al*, 2012; Virreira Winter *et al*, 2011). GRA15 also likely explains the difference in IRG coating of vacuoles of different type I strains. For example, the GT1 type I strain has a functional GRA15 while the RH type I strains do not. We previously showed that GT1 GRA15 can activate the NF-κB pathway and that vacuoles of the GT1 strain are significantly more coated with IRGs compared to RH vacuoles (Yang *et al*, 2013). Although both RH and GT1 have a lethal dose of just a single parasite in most laboratory mouse strains, deletion of *ROP18* makes GT1 avirulent after low dose infection (Shen *et al*, 2014) while deletion of *ROP18* in RH only delays death of the mice (Alaganan *et al*, 2014; Shen *et al*, 2014). It is likely that this difference is mediated by GRA15 as the presence of GRA15 in GT1 would make it more susceptible to IRGB6 PVM coating and subsequent parasite elimination.

Other studies have shown that IFNγ-dependent induction of IDO expression plays a role in inhibiting *Toxoplasma* growth in human fibroblasts (Bando *et al*, 2018; Pfefferkorn, 1984; Pfefferkorn *et al*, 1986). However, we find that inhibition of parasite growth is minimally dependent on IFNγ-dependent L-Trp breakdown (this study and Niedelman et al., 2013) (Niedelman *et al*, 2013). These differences might be due to fibroblasts derived from different tissues and/or the use of primary *vs*. transformed fibroblast (Bando *et al*, 2018), as the origin of the fibroblasts in other studies is unclear (Pfefferkorn, 1984; Pfefferkorn *et al*, 1986).

Thus, although we previously thought that the main role of GRA15 was the activation of the NF-κB pathway, it is possible that its primary role is reducing parasite virulence by enhancing IFNγ-mediated vacuole destruction. Furthermore, it was recently shown that GRA15 promotes the type I interferon response in mice by mediating STING polyubiquitination and enhanced cGAS/STING signaling through its ability to bind TRAF molecules (Wang *et al*, 2019a). Taken together, by observing the detrimental effect of GRA15 on the parasite, it might seem disadvantageous for *Toxoplasma* to have GRA15. However, the rapidly replicating tachyzoite *Toxoplasma* stages present during acute infection are not orally infectious, in contrast to the slowly replicating encysted bradyzoite stages. Therefore, to enhance its chances of transmission, *Toxoplasma* needs to balance immune evasion, to enable replication and dissemination, and immune activation, to prevent killing its host before orally infectious tissue cysts are formed. The contrasting goals of immune evasion and immune activation are reflected in *Toxoplasma*’s arsenal of secreted effectors. GRA15 is clearly an effector that makes the parasite less virulent and helps the host survive. In contrast to GRA15, multiple *Toxoplasma* effectors mediating resistance against IFNγ-mediated toxoplasmacidal mechanisms in murine cells have been identified (Hakimi *et al*, 2017; Hunter & Sibley, 2012). For example, ROP5 and ROP18, together with ROP17 and GRA7, cooperatively inactive the IRGs and thereby enhance *Toxoplasma* virulence in mice (Alaganan *et al*, 2014; Etheridge *et al*, 2014). However, type III strains and certain atypical strains (P89 and CASTELLS) do not express ROP18 because of a large insertion in their *ROP18* promoter region, while the BOF strain has very low expression of *ROP5* (Niedelman *et al*, 2012). Similarly, *Toxoplasma* secretes the effector IST beyond the vacuole and IST inhibits STAT1 transcriptional activity thereby enhancing *Toxoplasma* virulence (Gay *et al*, 2016; Olias *et al*, 2016). However, it also secretes the GRA24 effector beyond the vacuole which activates P38 MAPK thereby activating host immune responses (Braun *et al*, 2013). The ultimate outcome of infection is therefore determined by the exact combination of these, often polymorphic, effectors and the species or exact genetic background of the host. A combination of effectors that is optimal in one host might kill another host or lead to complete elimination of parasites in yet another host. It is these evolutionary forces that have likely led to differences in GRA15 expression level and/or sequence in different *Toxoplasma* strains.

In this manuscript, we show how the polymorphic effector GRA15 determines the differential susceptibility of type I RH and type II Pru strains to IFNγ-mediated growth inhibition in human and murine cells. This will help to understand the molecular basis of pathogenicity of different *Toxoplasma* strains in humans.

## Materials and Methods

### Reagents and antibodies

All the reagents, antibodies, primers, gDNAs and siRNAs used in this study are described in detail in **Suppl. Table 1**.

### Culture of cells and parasites

Human foreskin fibroblasts (HFFs) were routinely maintained in DMEM with high glucose (Gibco, Invitrogen) supplemented with 10% FBS, L-Glutamine (2mM), 100 U/mL Penicillin, 100 μg/mL Streptomycin and 20 μg/mL Gentamycin (complete medium). HFFs were passed using 0.25% trypsin. For all the experiments, HFFs were used at passage 5-10, but for serial passage of the parasites, higher passage number of the HFFs were used. Mouse embryonic fibroblasts (MEFs) were maintained in the complete medium supplemented with 10 mM HEPES, 1 mM sodium pyruvate and 1× MEM nonessential amino acids. MEFs were passed using 0.05% trypsin-EDTA. NF-κB *p65*^−/−^ MEFs were a gift from A. Sinai (University of Kentucky College of Medicine, Lexington, KY), and *Traf6*^−/−^ MEFs were provided by K. Fitzgerald (University of Massachusetts Medical School, Worcester, MA). All parasite lines were maintained *in vitro* by serial passage on monolayers of HFFs and cultured in DMEM with high glucose medium supplemented with 1% FBS, L-Glutamine (2mM), 100 U/mL Penicillin, 100 μg/mL Streptomycin and 20 μg/mL Gentamycin. A *Toxoplasma gondii* RH (RH 1-1) strain expressing click beetle luciferase and green fluorescence protein (GFP) and a Pru strain expressing firefly luciferase and GFP (Pru∆*hpt*, PruA7) were used as representative of type I and type II, respectively (Boyle *et al*, 2007). A genetically engineered RH strain expressing type II GRA15, Pru∆*gra15* and Pru∆*gra15* + GRA15-HA (GRA15 complemented strain) were described previously (Rosowski *et al*, 2011).

### Generation of TRAF6 knockdown human primary foreskin fibroblasts

HFFs were seeded in 12 well plates at a concentration of 2 × 10^5^ cells in 2 ml of complete media mentioned above for culturing HFFs, the night before the transduction. The next day, when cells were at ~70% confluency media were replaced with fresh complete media supplemented with polybrene (8 μg/ml, EMD Millipore). Ready to use lentivirus particles containing 4 different siRNAs targeting TRAF6 (**Suppl. Table 1**) or containing scrambled siRNA (**Suppl. Table 1**) were then used for transfection using 3 different MOIs (2/5/10) in 3 different wells for each virus particles (abm Inc, BC, Canada). Cells were then kept at 37°C, 5%CO_2_ inside an incubator overnight. The next day, cells were checked under an inverted fluorescence microscope for expression of GFP as the lentivirus expresses GFP as a fusion protein with the puromycin resistance gene and kept for an additional 24 h in the incubator. Following this, media was replaced with fresh media with puromycin (1.5 μg/ml) for 72 h for selection of the stably transduced cells, at this time puromycin was able to kill all the cells in untransduced cells, seeded in parallel in the same plate. Repeating the selection one more time for another 48 h cells were divided in 6 well plates to check by immunoblot for TRAF6 expression and knockdown of TRAF6 was confirmed.

### Generation of *MYR1* ko parasite

To generate the *MYR*1 insertional mutant in the Pru*Δku80* strain, the parasites were co-transfected with a mixture of the pTOXO_Cas9CRISPR:sg*MYR*1 vector with purified amplicons containing the *DHFR* cassette flanked by sequences homologous to the sequence targeted by sgMYR1 (5:1 mass ratio). These amplicons were generated by PCR amplification of the DHFR cassette using the primers *MYR*1-DHFR - F and *MYR*1-DHFR – R, and a vector carrying the *DHFR* cassette as template (Donald & Roos, 1993). Stable recombinants were selected with 1 μM pyrimethamine, single cell cloned by limiting dilution and verified by PCR analysis.

### GRA15 TRAF-binding site mutant generation

GRA15 TRAF2/6 binding site mutation constructs were amplified from pTKO-att-GRA15_||_HA vector (Rosowski *et al*, 2011) using specific primers (see primers Suppl. Table 1) and confirmed by sequencing. 50 μg of circular vectors (pTKO-att-GRA15_||_HA, pTKO-att-GRA15_||_-TRA2Fmut-HA and pTKO-att-GRA15_||_-TRAF2/6mut-HA) were transfected into 1 ×10^7^ RH∆*hxgprt* parasites by electroporation. Stable integrants were selected in media with 50 μg/ml mycophenolic acid and 50 μg/ml xanthine and cloned by limiting dilution. Expression and correct localization GRA15_||_ to the PVM were confirmed by IFA for HA and Western blotting.

### *In vitro Toxoplasma* infection

Parasites for *in vitro* infection were obtained from sequential syringe lysis using 27G and 30G needles of heavily infected HFF monolayers followed with a spin at 570 × g for 7 minutes. For the infection with RH strains MOIs of 1-3 and for Pru MOIs of 3-7 were used. Because RH and Pru strains often differ in viability and infectivity, equivalent ‘real’ MOIs were matched from plaque assay results performed for each experiment to be able to make strain comparisons. Following infection with *Toxoplasma*, each time plates were centrifuged at 160 × g for 3 minutes to synchronize the infection, prior to incubation for the required time.

### IFNγ stimulation of cells

In most of the experiments, HFFs were stimulated for 18-24 h in complete medium at 37 °C with 10 U/mL of human IFNγ (AbD Serotech, stock concentration is 10,000 U/mL). For some experiments, human IFNγ was used at concentration of 5-100 U/mL. MEFs were also stimulated for 18-24 h in complete medium with HEPES at 37 °C with 100 U/mL murine IFNγ (Peprotech, stock concentration is 100,000 U/mL).

### Use of inhibitors

For experiments using BAY11-7082 (1μM) or PYR41 (1 μM) or C25-140 (50 μM), those compounds were added 24 h post-stimulation with IFNγ but 2 h prior to infection and were kept throughout the infection for BAY11-7082 and C25-140 but washed away just prior to infection in the case of PYR41, as it is toxic to cells after longer incubation times. Bafilomycin A1 (100 nM) was added 1 h post infection as it affected the parasite invasion process.

### Luciferase assay for parasite growth

Luciferase assays were performed from the 96 well plates and for each strain and condition, triplicate wells were used. To the confluent monolayers of HFFs/MEFs (2 × 10^5^ cells/well), IFNγ was added (10 U/mL of human IFNγ and 100 U/mL of murine IFNγ) for 24 h prior to infection. Next day, infection with indicated parasites strains was done. For each strain 3 different MOIs were used for matching of results from similar parasite infectivity between the strains later with the plaque assay. For some experiment’s inhibitors were added either 2 h prior to infection or 1 h post infection as indicated. Following another 24 h incubation, culture supernatants were removed and 1× lysis buffer was added (Luciferase assay system, Promega) to the cells in the wells, followed by three freeze-thaw cycles. After that, 1× assay buffer containing luciferin was added to each well and transferred to clear centrifuge tubes for measurement of luciferase activity from the lysate using a single channel luminometer (Turner Biosystems). Luciferase reading of wells not treated with IFNγ was considered as 100% and relative growth was calculated for IFNγ- and inhibitor-treated wells.

### Immunofluorescence assays for recruitment of host markers to the PVM and percentage of infection

HFFs or MEFs were plated on coverslips in 24 well plates (1×10^5^ cells/well) and cultured, stimulated with IFNγ for 18-24 h, and subsequently infected with *Toxoplasma* for 24 h (to determine the percentage of infected cells and nuclear translocation of NF-κB), 3 h (to assess the recruitment of ubiquitin, p62, LC3B, GABARAP, GBP2, LAMP1 and TRAF6 in HFFs and MEFs) or 1 h (to determine IRGB6 coating in MEFs). Following incubation, cells were fixed with either 3% formaldehyde or 100% methanol depending on host marker (**See Suppl. Table 1**) and then permeabilized and blocked with either buffer containing 0.2% Triton X-100 along with 3% BSA and 5% goat serum (**See Suppl. Table 1**) or with buffer containing 0.2% freshly prepared saponin instead of Triton X-100 (**See Suppl. Table 1**). Cells were then treated with primary antibodies (**Suppl. Table 1**) for overnight incubation at 4 °C. Following that, each well was washed 3 times with 1x PBS and then secondary antibodies were added with Hoechst 33258 for 1 hr. Finally, coverslips were washed 5 times with 1× PBS and were mounted with VECTASHIELD antifade mounting medium. Imaging was done as described previously (Niedelman *et al*, 2013). For determination of nuclear translocation of NF-κB, nuclear intensity of at least 15 infected cells were taken into consideration, whereas to assess percentage of infection after 24 h, cells were counted in at least 6 independent fields and the values observed in untreated infected cells were taken as 100% and calculation for the rest was done relative to untreated infected cells. To measure the recruitment of host markers on the PVM, at least 100 infected cells were counted.

### Plaque assay

For the plaque assay, freshly confluent 24 well plates of HFFs or MEFs were used. The day before infection fresh media was added replacing the media from the plates and stimulated with 10 U/mL human IFNγ or 100 U/mL mouse IFNγ or left unstimulated for 24 h. For infection, freshly harvested parasites, 100 parasites of RH and 250 parasites of Pru strains were added to the 24 well plates. Infected plates were incubated for 4 days at 37 °C for RH strains and for 6 days in the case of Pru strains. For calculating the percentage of plaque loss, the following formula was used as described previously (Niedelman *et al*, 2012, 2013): [(Number of plaques in unstimulated condition - Number of plaques in stimulated condition)/Number of plaques in unstimulated condition] × 100. Plaque areas were captured and analyzed using a Nikon TE2000 inverted microscope equipped with Hamamatsu ORCA-ER digital camera, and NIS Elements Imaging Software, respectively. Plaque area loss was calculated using the same formula for plaque loss except using the plaque areas in place of plaque numbers. For all experiments, at least 20-25 plaques from technical duplicate wells were imaged.

### IDO activity assay

The IDO activity upon IFNγ stimulation and *Toxoplasma* infection was evaluated by measuring L-Kynurenine from the culture supernatant of HFFs. Cells were cultured in 96-well plates as mentioned in earlier assays with the complete DMEM medium containing total 0.6 mM L-Tryptophan (0.52 mM L-Tryptophan was added to existing 0.08 mM L-Tryptophan in the media). HFFs were either stimulated with 10 U/mL IFNγ for 18-24 h or left untreated and subsequently infected with parasite strains at different MOIs for 24 h before harvesting the culture supernatant. The concentration of L-Kynurenine was measured using 1.2% p-dimethylaminobenzaldehyde in glacial acetic acid solution (Ehrlich reagent). Briefly, 150 μL of culture supernatant was mixed with 20 μL of 30% trichloro acetic acid in a V-bottom 96 well plate followed with incubation at 50 °C for 30 minutes. Subsequently, the plate was centrifuged for 10 minutes at 600 × g, 100 μL of culture supernatant was mixed with Ehrlich reagent and incubated for 10 minutes and absorbance was recorded at 490 nm using a plate reader (Molecular Device Spctramax M2e). A standard curve of L-kynurenine (0-1500 μM) was used to calculate the concentrations in the samples.

### Co-Immunoprecipitation

40× T175 of human foreskin fibroblast (HFF) were stimulated or not with 10 U/ml of human IFN-γ (AbD Serotech) for 24 h. Then, 2 h before infection (MOI:5 to 10) with RH+GRA15_||_-HA or RH+GRA35-HA (for each parasites 10× T175 containing IFN-γ stimulated HFFs and 10 unstimulated) the cells were incubated with 50 μM of VX7655 (Selleck chem). 16 h after infection, the cells were washed once and scraped with cold PBS. The cells were centrifuged and resuspended in 6 ml of lysis buffer (HEPES 10mM pH7.9, MgCl2 1.5mM, KCl 10mM, EDTA 0.1mM, 0.5mM, NP40 0.65%, cocktail of protease inhibitor (Roche), phenylmethylsulfonylfluoride (PMSF) 0.5mM.) for 45 minutes at 4°C. 1% of these lysates were kept as input for the immunoblot. The lysate was centrifuged 30 minutes at 18,000 × g, 4°C. Each sample was incubated with 100 μl of HA magnetic beads (Thermo scientific) rotating overnight at 4°C. The beads were washed 3 times with Tris-HCl 10mM pH7.5, NaCl 150mM, Triton-100X 0.2%, PMSF 0.5mM, cocktail of protease inhibitor (Roche), once with Tris-HCl 62.5mM pH 6.8 and subsequently resuspended in 100 μl of this buffer performed immunoblotting (30 μl) and did the mass spec analysis from the rest of the samples.

To perform immunoprecipitation with GRA15 mutants in HFFs or in MEFs one T175 for each condition (untreated or IFNγ-treated) was infected with RH+GRA15_WT_-HA, RH+GRA15_TRAF2mut_-HA, RH+GRA15_TRAF2/6mut_-HA, or RH+GRA45-HA expressing parasites for 24 h in MEF and 8 h in HFFs. Following infection, cells were washed once with ice cold 1× PBS, and then scraped with ice cold PBS, centrifuged and resuspend in 1 ml of lysis buffer mentioned above for 30 minutes at 4°C. After that 10% of lysate was put aside for use as input during immunoblotting. The remaining lysates were then processed as mentioned above and incubated with 20 μl of HA magnetic beads overnight, in rotating condition at 4°C. Next day, beads were washed with the lysis buffer 3 times and subsequently resuspended in 60 μl of 1× loading dye to perform immunoblotting.

### Immunoblotting

20 μl of the HA magnetic beads of each sample was used to run on a 12% SDS-PAGE. The proteins were transferred to a PVDF membrane, blocked 30 minutes with TBST, 5% nonfat dry milk. The membrane was blotted overnight at 4°C with rat antibody against the HA tag (Roche, 1/500 dilution), TRAF2 and TRAF6 rabbit antibodies (**Suppl. Table 1**) followed by respective secondary HRP antibodies (**Suppl. Table 1**).

### Mass spectrometry-based proteomics

The HA magnetic beads were sent to the Proteomic Core Facility of University of California, Davis for mass spectrometry analysis. Briefly, the proteins were digested using Promega modified trypsin overnight at room temperature on a gently shaking device. Resulting peptides were analyzed by online LC-MS/MS Q-Exactive. All MS/MS samples were analyzed using X! Tandem (The GPM, thegpm.org; version X! Tandem Alanine (2017.2.1.4)). X! Tandem was set up to search the uniprotHSTG_crap database assuming the digestion enzyme trypsin. X! Tandem was searched with a fragment ion mass tolerance of 20 PPM and a parent ion tolerance of 20 PPM. Glu->pyro-Glu of the n-terminus, ammonia-loss of the n-terminus, gln->pyro-Glu of the n-terminus, deamidated of asparagine and glutamine, oxidation of methionine and tryptophan, dioxidation of methionine and tryptophan and dicarbamidomethyl of lysine were specified in X! Tandem as variable modifications. Scaffold (version Scaffold_4.8.6, Proteome Software Inc., Portland, OR) was used to validate MS/MS based peptide and protein identifications. Peptide identifications were accepted if they could be established at greater than 50.0% probability by the Scaffold Local FDR algorithm. Peptide identifications were also required to exceed specific database search engine thresholds and X! Tandem identifications were also required. Protein identifications were accepted if they could be established at greater than 9.0% probability to achieve an FDR less than 5.0% and contained at least 1 identified peptide. Protein probabilities were assigned by the Protein Prophet algorithm (Nesvizhskii *et al*, 2003). Proteins that contained similar peptides and could not be differentiated based on MS/MS analysis alone were grouped to satisfy the principles of parsimony. Proteins sharing significant peptide evidence were grouped into clusters.

### Statistical analysis

All statistical analyses were performed using Graph Pad prism version 7.0. All the data presented are mean ± standard error of mean (SEM) and the exact n values are mentioned in each of the figure legends. For all the calculations, p-values of <0.05 are considered as significant. Parameters with 2 different variables and groups were compared by Two-way ANOVA followed with either Bonferroni or Tukey’s multiple comparison test. Parameters with one variable but three or more groups were compared by One-way ANOVA followed with Tukey’s multiple comparison test. For one variable test with two groups, two-tailed unpaired t-test was used. Specific statistical test performed for each figure was stated in each of the figure legend.

## Data availability

The authors declare that all data supporting the findings of this study are available within the article and its Supplementary Information files or are available from the authors upon request. All unique materials (e.g. the diversity of parasite lines described in this manuscript) are available upon request (contact).

## Acknowledgment

We thank all members of the Saeij laboratory for productive discussions and Dr. Kevin Woolard for providing the instrumental facilities for assistance in the project. JPJS was supported by the National Institutes of Health-NIH-2R01AI080621-06A1. DM was supported by American Heart Association Post-doctoral fellowship (18POST34030036)

## Author contribution

DM and JPJS designed the study. DM and LOS performed and interpreted the experimental work. LB generated *MYR1* knockout parasite strains. MAH provided insightful discussions and constructive suggestions and supervised the generation of knockout parasite strains. JPJS supervised the research. DM and JPJS wrote the paper.

