## Supplementary Files for "*Toxoplasma* GRA15 limits parasite growth in IFNγ-activated fibroblasts through TRAF ubiquitin ligases"

**Supplementary information**

**Supplementary Figure 1**

HFFs were pre-stimulated for 24 h with 10U/mL IFNγ followed by infection with the indicated strains. **a)** Plaque assays were performed with PruΔ*ku80*Δ*hpt* (Pru), PruΔ*ku80*Δ*hpt*Δ*myr1* (PruΔ*myr1*) strains and plaque numbers were counted and plaque areas were measured after 6 days. Plaque numbers and plaque areas are averages from eight and six biological replicates, respectively. **b)** Cells were supplemented with media containing 0.6 mM L-Trp, stimulated with IFNγ for 24 h and infected with RH or Pru parasites. Plaque number was measured 4 (RH) or 6 days (Pru) p.i. Experiments were performed 2 times. **c)** L-Kynurenine was measured from HFF culture supernatant as a marker of IDO activity. Experiment was done 2 times. Statistical analysis was done by two sample student's t test (a). Data are represented as mean ± SEM.

**Supplementary Figure 2**

**a)** PVM ubiquitin coating intensity is similar between RH, Pru and PruΔ*gra*15. The measurement of fluorescence intensity of ubiquitin staining on the PVM was performed using NIS element software version 4 (Nikon) from the experiments described in figure 2. For each strain type intensities of at least 50 vacuoles was measured. **b)** p62 coating was performed as described in Figure 2, except here it is performed with RH, GT-1 and RH+GRA15$_{II}$. Experiment was performed 3 times with RH and RH+GRA15$_{II}$ and 2 times with GT-1. **c)** Plaque assay was performed as described in figure 1a. Plaques were counted 4 and 6 days p.i. for RH and GT-1, respectively. Experiment was performed three times. Statistical analysis was done by One-way ANOVA followed with Tukey's multiple comparison test for a and two sample student's t test for b-c. Data are represented as mean ± SEM.

**Supplementary Figure 3**

Panel of Pru-infected HFFs that were previously stimulated with 10U/ml of IFNγ shows different LAMP1 staining patterns characteristic of endo-lysosomal fusion and parasite degradation. Cells were stimulated, infected, fixed and stained as mentioned in figure 3. Yellow arrows indicated the LAMP1-coated parasites.

**Supplementary Figure 4**

HFFs in 24 well plates were stimulated with IFNγ (10 U/mL for 24 h) and subsequently treated with **a)** E2-ubiquitin conjugating enzyme inhibitor, BAY11-7082 (1 μM) for 2 h and subsequently infected with Pru parasites for 3 h. The percentage of vacuoles that stained positive for ubiquitin and p62 was scored. Experiment was performed three times. **b)** TRAF6 (E3) -ubiquitin ligase inhibitor, C25-140 (50 μM) for 2 h and subsequently infected with Pru parasites for 3 h. The percentage of vacuoles that stained positive p62 was determined. Experiment was performed three times. Statistical analysis was done by One-way ANOVA followed with Tukey's multiple comparison test for a and b. Data are represented as mean ± SEM. **c-e)** Complete Western blot pictures for figure 4c. Membrane was blotted for with antibodies against HA (c), TRAF2 (d) or TRAF6 (e). The red boxes indicate the portions of the blot that was cropped to show in the main figure. **f-h)** Complete Western blot pictures for figure 4e left panel. Membrane was blotted for with antibodies against HA (f), TRAF2 (g) or TRAF6 (h). The red boxes indicate the portions of the blot that was cropped to show in the main figure. Between the input lane and immunoprecipitate lane, one additional lane was not used due to spillover of input from the uninfected HFFs. **i-k)** Complete Western blot pictures for figure 4e right panel. Membrane was blotted for with antibodies against HA (i), TRAF2 (j) or TRAF6 (k). The red boxes indicate the portions of the blot that was cropped to show in the main figure. **l-m)** Complete Western blot pictures for figure 4j. Membrane was blotted for with antibodies against TRAF6 (l) and GAPDH (m). The red boxes indicate the portion of the blot that was cropped to show in the main figure.

**Supplementary Figure 5**

**a)** Identification of putative TRAF2 and TRAF6 binding sites in type II GRA15 protein. The amino acid sequence of type II GRA15 was derived from ToxoDB (https://toxodb.org/toxo/). TRAF2 binding sites are highlighted in red while TRAF6 binding site is highlighted in yellow (Sangaré *et al*, 2019). **b)** Ubiquitination sites within type II GRA15 sequence was identified using http://www.ubpred.org/ and the score provided was also derived from the same online tool. **c)** Identification of putative ubiquitinated p62/TRAF6 acceptor sites within type II GRA15 sequence as described by Jadhav et al. 2008 (Jadhav *et al*, 2008). **d)** Results from the DNA sequencing of the RH expressing GRA15 TRAF2 binding mutants or TRAF2/6 binding mutants. In the right column the nucleotide positions of the displayed sequence read are mentioned, in the middle column, the exact sequence was shown and in the right column, the corresponding changes in the amino acids are also mentioned. **e)** The localization of HA-tagged GRA15 in RH expressing either GRA15 wild-type, GRA15 TRAF2 binding mutant, or GRA15 TRAF2/6 binding mutants, shown using immunofluorescence using anti-HA antibody. **f)** The expression of GRA15 among the RH strains expressing either GRA15 wild-type, GRA15 TRAF2 binding mutant, or GRA15 TRAF2/6 binding mutants (upper panel). Expression was normalized to the GRA1 expression of the parasites and plotted (lower panel).

**Supplementary Figure 6**

**a-c)** Complete Western blot pictures for figure 6a. Membrane was blotted with antibodies against HA (a), TRAF2 (b) or TRAF6 (c). The red boxes indicate the portions of the blot that was cropped to show in the main figure. Between the input lane and immunoprecipitate lane, one additional lane was not used due to spillover from the lane before. **d-f)** Complete Western blot pictures for figure 6e. Membrane was blotted for with antibodies against HA (d), TRAF6 (e) or TRAF2 (f). The red boxes indicate the portions of the blot that were cropped to show in the main figure.

#### Supplementary Figure 1

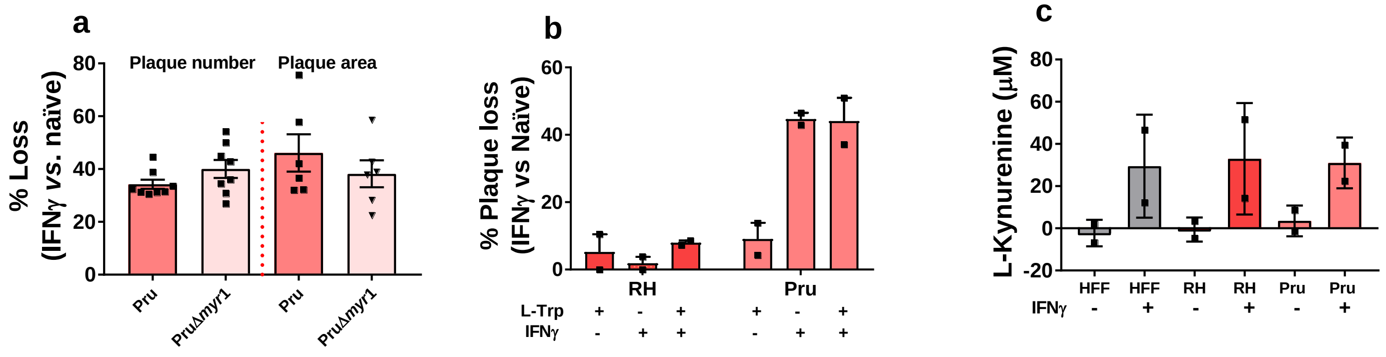


#### Supplementary Figure 2

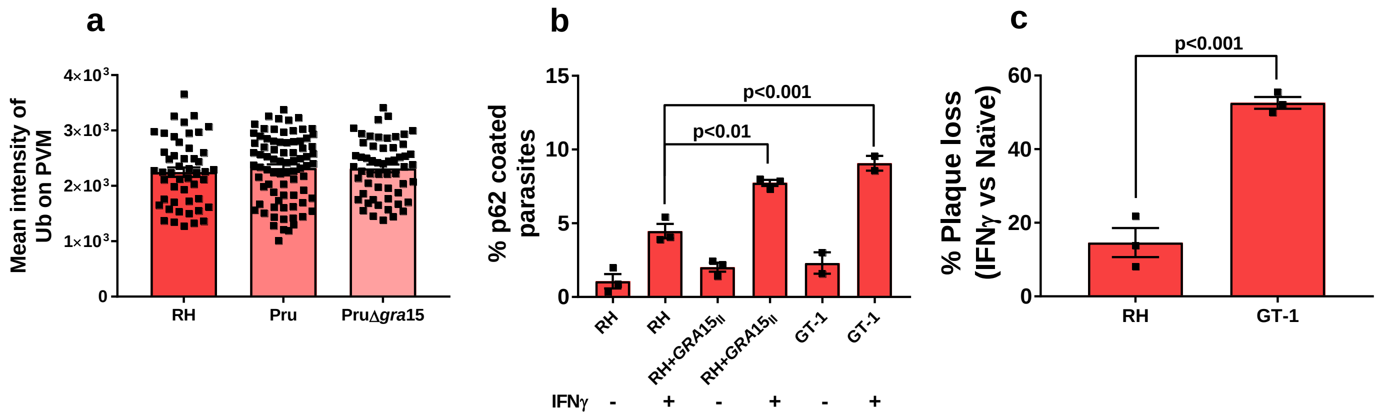

Supplementary Figure 3

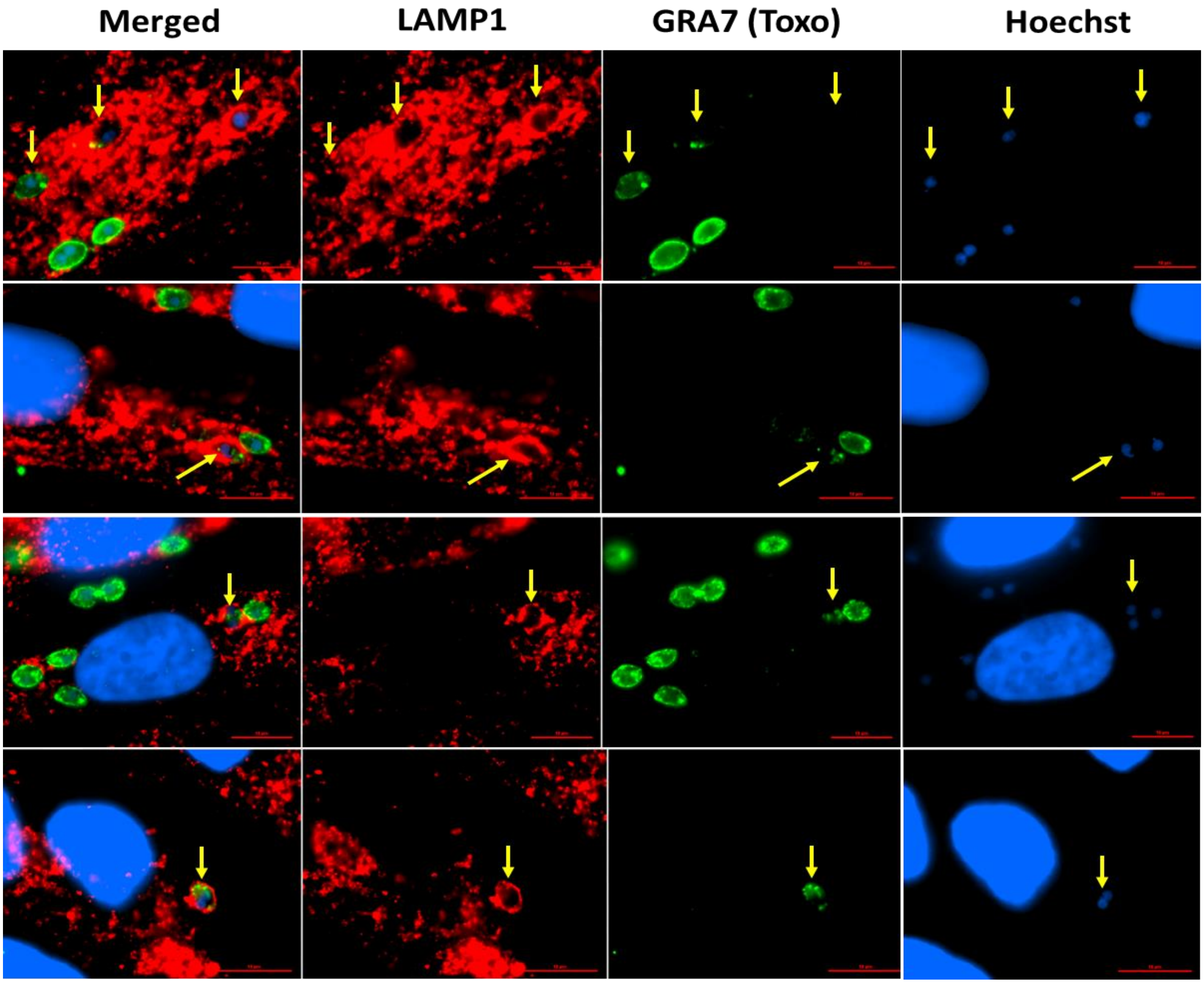

### Supplementary Figure 4

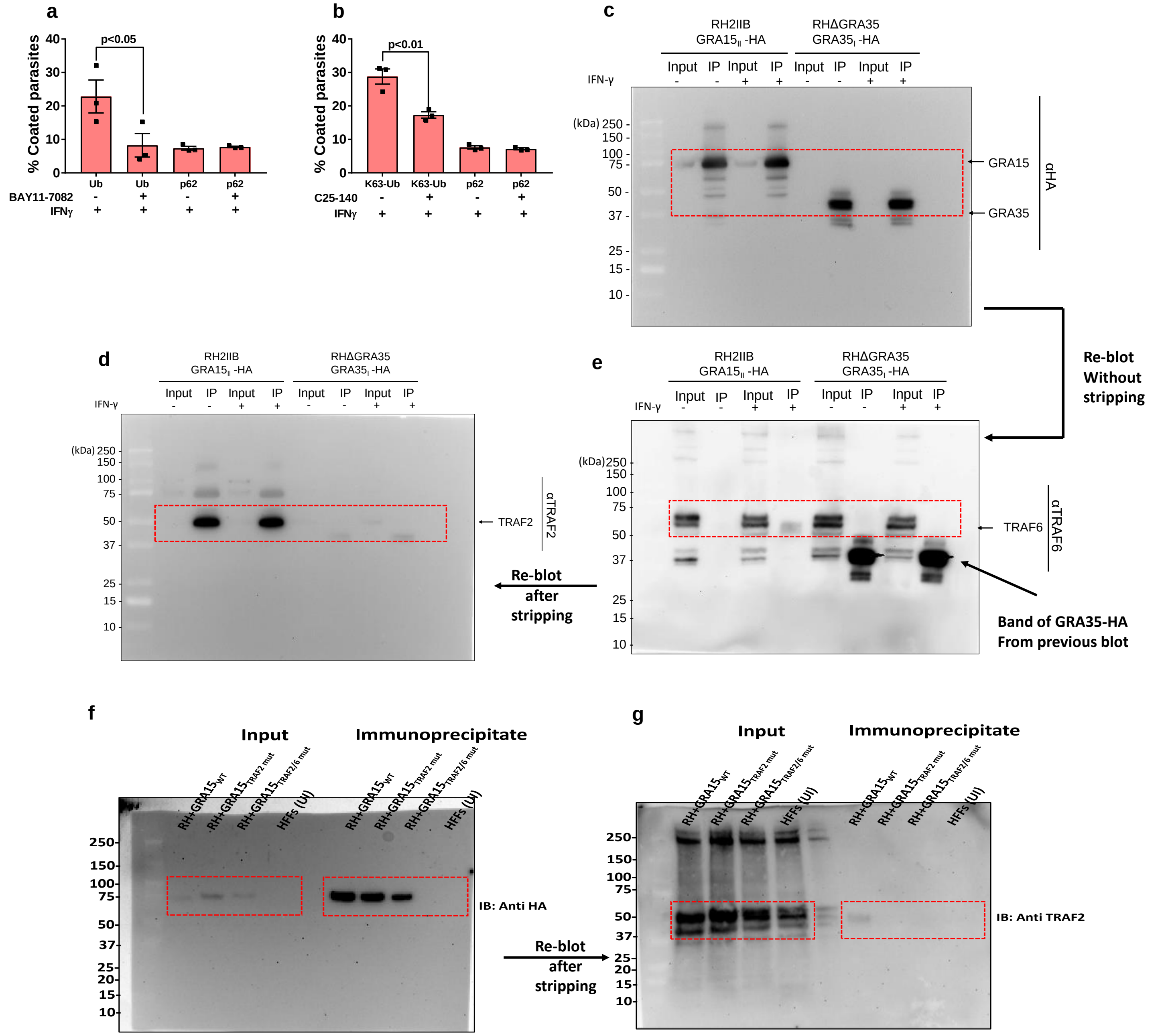

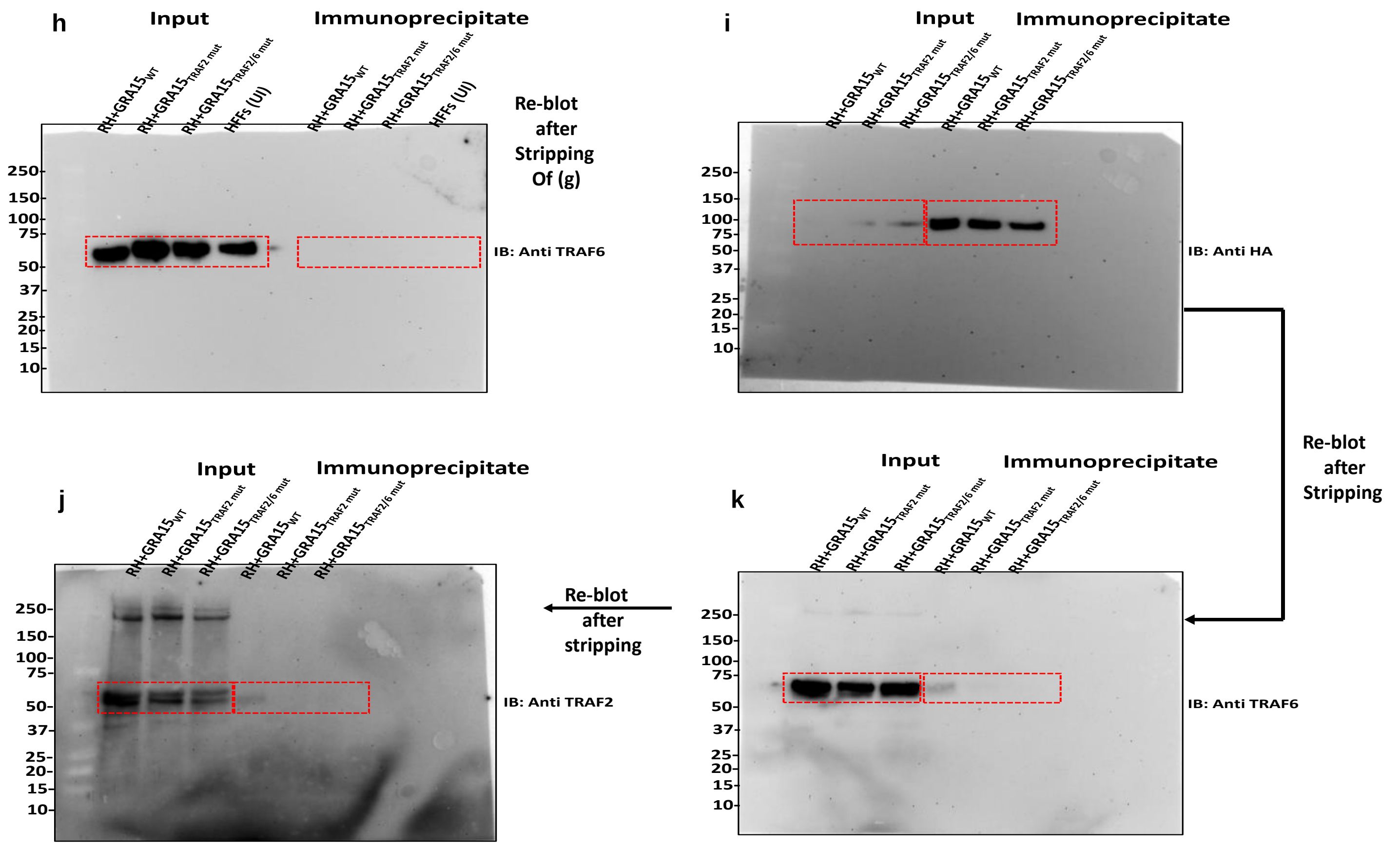
h
Input
Immunoprecipitate
RH+GRA15WT
RH+GRA15TRAF2 mut
RH+GRA15TRAF2/6 mut
HFFs (UI)
Re-blot after Stripping Of (g)
IB: Anti TRAF6
i
IB: Anti HA
Re-blot after Stripping
j
IB: Anti TRAF2
Re-blot after stripping
k
IB: Anti TRAF6

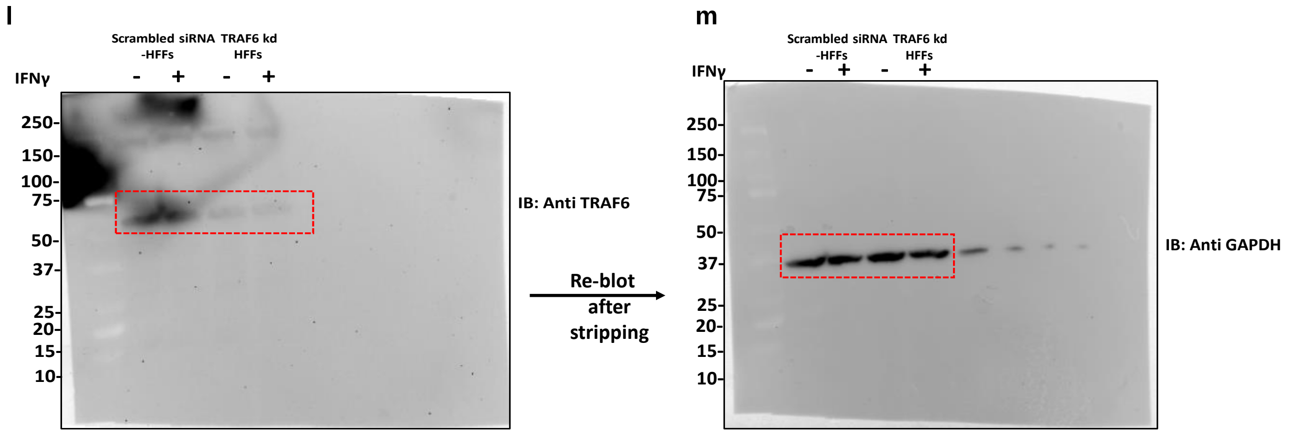
l
Scrambled siRNA -HFFs
TRAF6 kd HFFs
IFNγ
IB: Anti TRAF6
Re-blot after stripping
m
IB: Anti GAPDH

### Supplementary Figure 5

#### a. Identification of TRAF2 and TRAF6 sites in GRA15

>GRA15_Type II (550 amino acids)

```
MVTTTTPTPPGAPAVVPIFDVVYQLNPHVFRSRFSRRNRARRVASSKSRSIIRWLGYLTVLAAVILLGAY
AVRRVSRDLSDSVRETRRGRRITGSVPPGTTRPRSESCTGTQVDGGCGADTSTDGKSESEQTENGEDSRF
STRTPIHVTASTSPFATRKAAEERSSSPRDRKVPEGAQLPTSSTPHAQRKDSGSDSRNPSTLIPSPGTNT
FNMNFYIIGAGSSALDFIFPHTPDAQATVVSPPRSAAAAPTVETVPRVRTYSTPTTLTLPTAPATATSNH
MHASATPSPPERPQNFRGGLMRQNGMVEGTSLTTTEAGMPAPLQSPQHIETEARLTYSNHLKSPHTPETP
TVHSIDPVVGTSGHSVAVGSQSPAGGPPTDSRTPAALTPTSSSFSHADSLETSEHPQSGPSLHPLISGIQ
DAVQSQLPLSQQETLPVVENATFFGPQQTPPWMDETAAAAIPLAPSQPGSRTQPISSPHTLLSRSGGVSA
VPGPPRTENPRQPQVPGENSYYSVPTERTISDPGPQRAGAAQADGIGAGGPRDTQSAVTP
```

***TRAF2 binding site = (P/S/A/T)X(Q/E)E (X=any amino acid)**
***TRAF6 binding site = PXEXXZ (Pro/any amino acid/Glu/any amino acids/any aromatic and acidic amino acid)**

#### b. Identification of ubiquitination sites in GRA15

>GRA15_Type II (550 amino acids)

```
MVTTTTPTPPGAPAVVPIFDVVYQLNPHVFRSRFSRRNRARRVASSKSRSIIRWLGYLTVLAAVILLGAY
AVRRVSRDLSDSVRETRRGRRITGSVPPGTTRPRSESCTGTQVDGGCGADTSTDGKSESEQTENGEDSRF
STRTPIHVTASTSPFATRKAAEERSSSPRDRKVPEGAQLPTSSTPHAQRKDSGSDSRNPSTLIPSPGTNT
FNMNFYIIGAGSSALDFIFPHTPDAQATVVSPPRSAAAAPTVETVPRVRTYSTPTTLTLPTAPATATSNH
MHASATPSPPERPQNFRGGLMRQNGMVEGTSLTTTEAGMPAPLQSPQHIETEARLTYSNHLKSPHTPETP
TVHSIDPVVGTSGHSVAVGSQSPAGGPPTDSRTPAALTPTSSSFSHADSLETSEHPQSGPSLHPLISGIQ
DAVQSQLPLSQQETLPVVENATFFGPQQTPPWMDETAAAAIPLAPSQPGSRTQPISSPHTLLSRSGGVSA
VPGPPRTENPRQPQVPGENSYYSVPTERTISDPGPQRAGAAQADGIGAGGPRDTQSAVTP
```

**Output:**

| Residue | Score | Ubiquitinated |
|---|---|---|
| 47 | 0.32 | No |
| 126 | 0.95 | Yes High confidence |
| 159 | 0.81 | Yes Medium confidence |
| 172 | 0.95 | Yes High confidence |
| 190 | 0.94 | Yes High confidence |
| 342 | 0.89 | Yes High confidence |

| Label | Score range | Sensitivity | Specificity |
|---|---|---|---|
| Low confidence | 0.62 = s = 0.69 | 0.464 | 0.903 |
| Medium confidence | 0.69 = s = 0.84 | 0.346 | 0.950 |
| High confidence | 0.84 = s = 1.00 | 0.197 | 0.989 |

#### c. Sequence similarities flanking the TRAF6/p62 ubiquitin acceptor sites within GRA15

| Consensus pattern | * | K | * | X | X | * | ! | * | !! | * |
|---|---|---|---|---|---|---|---|---|---|---|
| Site 125-134 (K126) | G | K | S | E | S | E | Q | T | E | N |
| Site 158-167 (K159) | R | K | A | A | E | E | R | S | S | S |
| Site 171-180 (K172) | R | K | V | P | E | G | A | Q | L | P |
| Site 189-198 (K190) | R | K | D | S | G | S | D | S | R | N |
| Site 341-350 (K342) | L | K | S | P | H | T | E | T | P | T |

(*) any hydrophobic amino acids; (!) polar amino acid 1 (Q/Y/C/S); (!!) polar amino acid 2 (H/D/T); (x) any amino acids.

#Amino acids in Red in the putative ubiquitination sites share similarities with the TRAF6/p62 consensus sequences

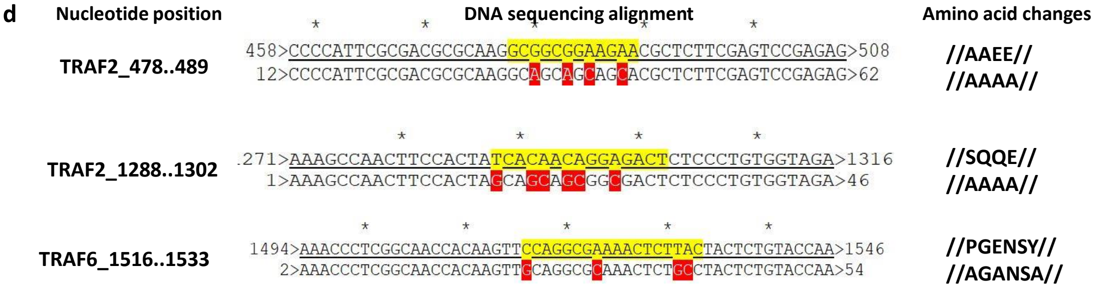


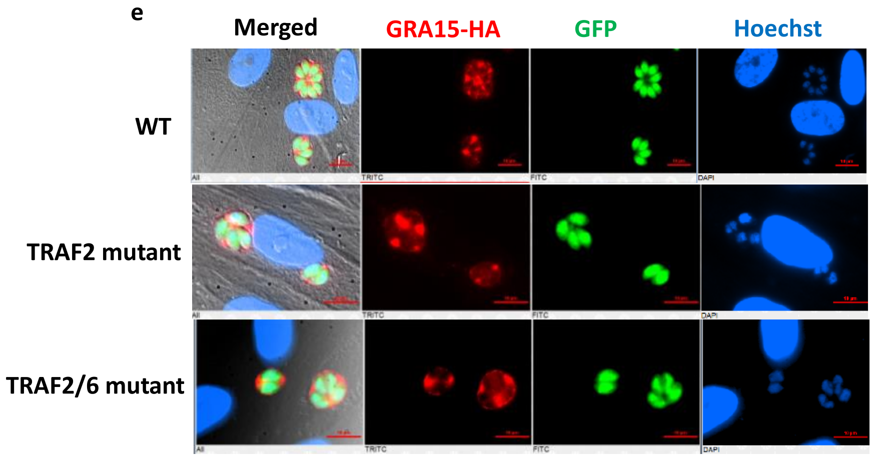


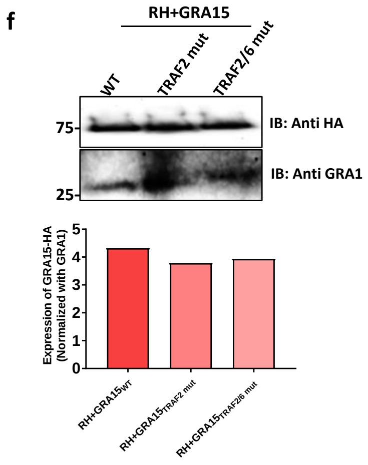

### Supplementary Figure 6

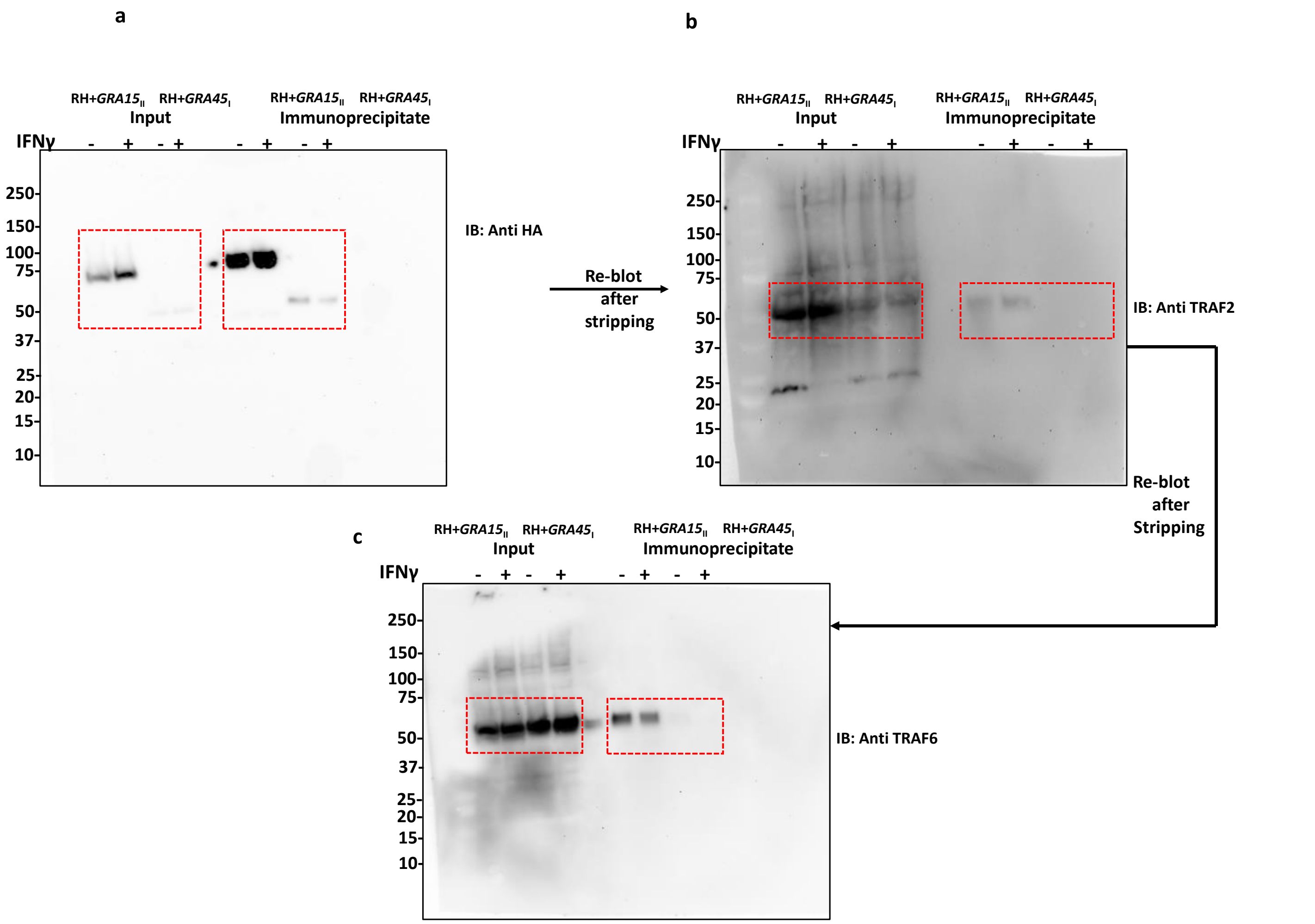

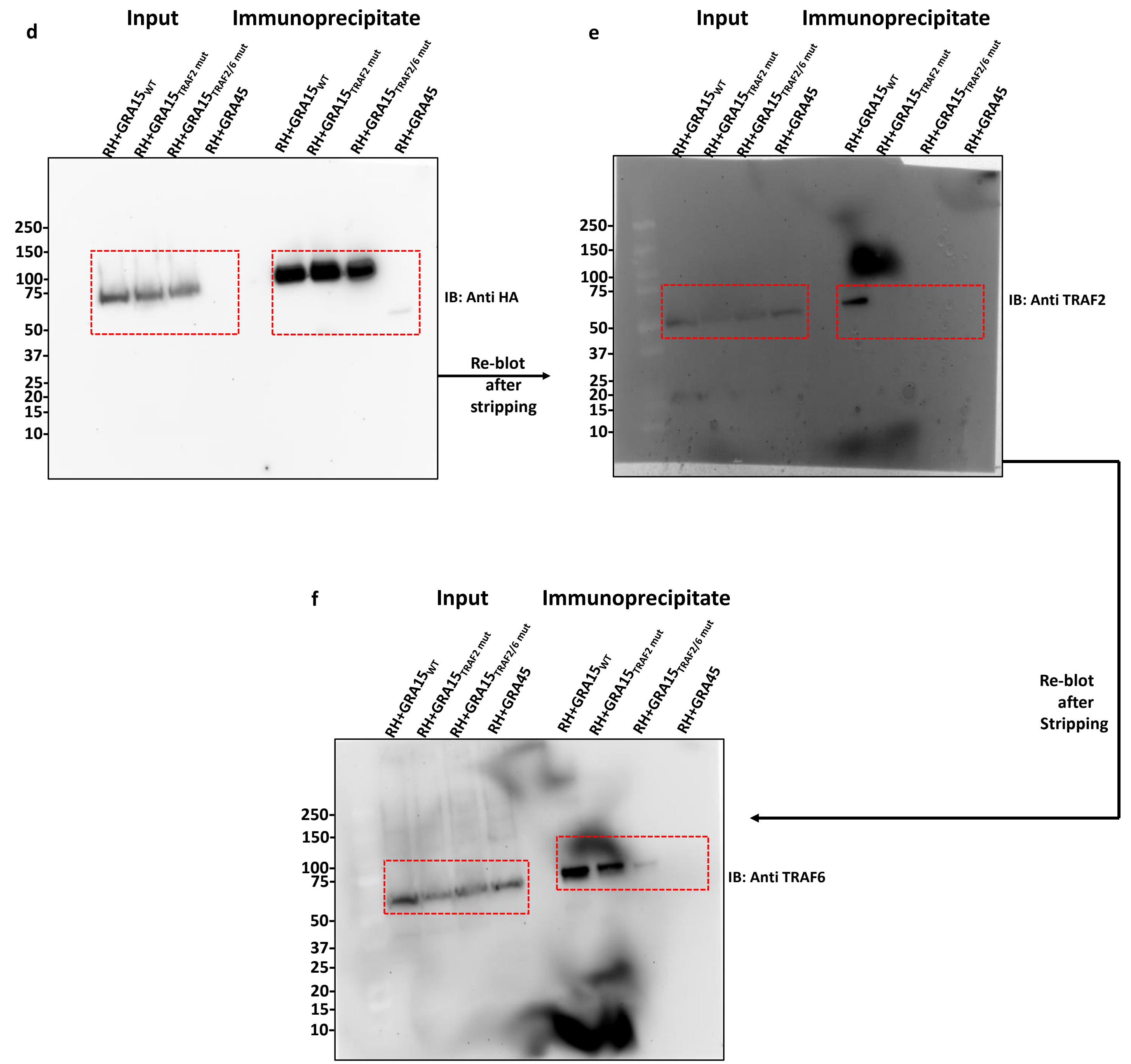

Activates the NF-κB Pathway through Interactions with TNF Receptor-Associated Factors. *MBio* **10:** e00808-19 Available at: http://mbio.asm.org/content/10/4/e00808-19.abstract
